# Atomic layer deposition for core-shell microparticle vaccines enabling programmable antigen delivery and enhanced humoral immune responses

**DOI:** 10.64898/2026.05.29.728600

**Authors:** Namit Chaudhary, Holly J. Coleman, Elias V. Paolone, Jiancheng Yu, Amir W. Ledbetter, Ashley A. Lemnios, Agnes A. Walsh, Heikyung Suh, Alexander Wang, Maryam M. Mansoor, JoLynn B. Giancola, Mariane B. Melo, Amber M. Rauch, Erika S. Langsfeld, Urvi R. Parlikar, Marisa O. Pacheco, Carly A. Williams, Hans H. Funke, Robert L. Garcea, Theodore W. Randolph, Darrell J. Irvine

## Abstract

Technologies that simplify complex dosing regimens as single-shot immunizations may be important for vaccines against difficult to neutralize pathogens. Here we characterized delivery mechanisms of core-shell microparticles comprising clinically-relevant HIV immunogen formulations spray-dried to form solid spherical microparticles and subsequently coated with a nanoscopic alumina shell using atomic layer deposition (ALD). ALD particles exhibited a time delay in antigen release programmed by the alumina shell followed by prolonged antigen release, which steadily accumulated in antigen-presenting cells at the injection site and draining lymph nodes and accumulated on follicular dendritic cells in B cell follicles. ALD vaccines elicited continuous expansion of antigen-specific germinal center B cells over 8 weeks, serum antibody responses with >10-fold slower antigen-binding off-rates, and 2-fold more long-lived plasma cells compared to traditional bolus vaccination with potent adjuvants. Single administration ALD technology thus promotes key events in the primary immune response important for vaccines against HIV and other challenging pathogens.

## INTRODUCTION

Successful vaccines targeting infectious disease-causing pathogens have saved millions of lives over the past century (*1, 2*). Most vaccines are thought to function through the induction of protective antibody responses. These develop when antigen-specific B cells enter germinal centers (GCs) and undergo the cyclic process of proliferation and somatic hypermutation of their antibody genes in a Darwinian selection process, producing a diversified pool of antibodies that bind with high affinity to the target antigen (*3*). Many vaccines elicit protective immunity following two or three bolus administrations, which in some cases can be quite durable over time (*4*). However, there are several pathogens for which highly effective vaccine development has remained elusive, including HIV, malaria, and tuberculosis. In addition, there is great interest in the development of vaccines that could broadly protect against pathogen variants or classes of pathogens. Examples in this category include “universal” influenza vaccines, pan-sarbecovirus vaccines, or pan-flavivirus vaccines (*5–7*). In principle, vaccines should be possible for many of these pathogens/scenarios because broadly neutralizing antibodies (bnAbs, e.g., antibodies that can neutralize diverse strains of the pathogen of interest) have been identified from infected humans for many of these diseases (*6, 8, 9*). Thus, a central challenge in vaccine development is to understand how to reliably elicit such bnAb responses by vaccination.

A common feature of the outstanding vaccine challenges noted above is that protective B cells in the naïve human repertoire are extremely rare (1 in >10^6^ naïve B cells in some scenarios (*10, 11*)), and generation of bnAbs from these naïve B cells requires a high level of somatic hypermutation from the naïve germline antibody sequence. Taking the case of HIV as the prototypical exemplar of this challenge, many efforts in the field are focused on vaccination strategies that employ engineered priming antigens (“germline targeting” immunogens) designed to effectively activate bnAb precursor B cells, followed by a series of booster immunizations aiming to shepherd mutations in the responding B cell population to eventually reach the production of protective antibodies (*12, 13*). Preclinical studies suggest that vaccine regimens comprising as many as 5 or 6 boosts may be required to induce bnAbs to some sites of vulnerability on HIV, vaccine regimens that would be a major challenge to implement for global vaccination campaigns (*14–18*). In addition, to most effectively engage protective precursor B cells, “extended dosing” regimens may be important, which prolong antigen exposure to the immune system and lead to amplified and prolonged GC responses that allow rare and subdominant B cell clones to be activated and expanded. Proof of concept of the beneficial effects of extended-dosing vaccination in HIV has to date been demonstrated using repeated-injection regimens (administering up to 7 doses over a 2-week period) or through implantation of osmotic pump devices for prolonged vaccine delivery (*19–22*). However, such approaches are also problematic for clinical vaccines.

An important class of materials that may be able to address these practical and immunological challenges together are controlled-release microparticles. Microspheres composed of biodegradable materials that either release encapsulated vaccines over a prolonged time period (*23–26*) or release vaccines after a programmed delay period (*27*) (for single shot prime-boost immunization) were first described in the late 1980s/1990s and have been heavily explored to promote protective immune responses. Such microparticles can co-encapsulate adjuvants or exhibit intrinsic adjuvant properties (*23, 25, 28*). While promising in preclinical studies, these early formulations faced challenges in manufacturing and retention of antigen integrity that hindered clinical translation (*29, 30*). Recently, a number of advances have been made that are renewing interest in controlled-release microparticles, including the development of methods to encapsulate vaccines in controlled-release microspheres under very mild conditions (*31, 32*), formulation strategies to protect antigens during encapsulation (*33, 34*), and new manufacturing approaches such as the use of microfluidic fabrication (*35*).

We recently reported a novel delivery technology based on the encapsulation of vaccines in engineered core-shell particles designed to provide programmed patterns of vaccine release *in vivo* (*36–40*). In this approach, liquid vaccines formulated with polysaccharides are spray-dried to embed and stabilize the vaccines within amorphous solid microparticle powders. The surfaces of these powders are then coated by atomic layer deposition (ALD), wherein alternating exposure to trimethyl aluminum vapor and water vapor leads to sequential, self-limiting chemical reactions at the particle surfaces that deposit single molecular layers of aluminum oxide (alumina) around the particle core. Application of repeated cycles of the ALD process forms multi-molecular, nanoscopic alumina shells of precisely defined, arbitrary thickness. These alumina shells form an impermeable barrier that prevents vaccine release from the particles until the alumina shell dissolves, allowing precise control over the timing of antigen release *in vitro* and immune response induction *in vivo* (*40–42*). Mixtures of ALD particles with thin alumina shells and particles with thicker shells thus have the potential to deliver prime and booster immunizations from a single injection. However, mechanisms of action underlying ALD vaccines remain poorly understood: First, the degree to which ALD encapsulation preserves antigen integrity has not been analyzed to date. Second, the pharmacokinetics and mechanisms of antigen delivery from ALD-coated particles to cells at the injection site and lymphoid organs are unknown. Finally, ALD subunit vaccines have been shown in some cases to elicit more potent antibody responses than unformulated control immunizations or antigens administered with traditional aluminum hydroxide or squalene-based adjuvants (*38, 40, 43*), but effects on the germinal center response and antibody affinity maturation of critical importance for pathogens such as HIV have not been assessed.

Here, using vaccines comprising stabilized HIV Env trimer immunogens, we characterized interactions of ALD particles with antigen presenting cells (APCs) *in vitro* and *in vivo*, analyzed the *in vivo* pharmacokinetics of ALD vaccines, and determined how ALD delivery alters the magnitude, character, and kinetics of humoral immune responses in mice. We found that ALD vaccines exhibit a multi-phase pattern of *in vivo* antigen pharmacokinetics, with an initial delay in antigen clearance from the injection site that is determined by the alumina shell thickness, followed by sustained, slow clearance of antigen away from the injection site over several weeks. In contrast to traditional bolus vaccine formulations characterized by rapid antigen clearance and low amounts of antigen capture by APCs, ALD vaccines elicit sustained accumulation of antigen in APCs at both the injection site and draining lymph nodes over at least three weeks once vaccine release initiates from ALD particles. In parallel, antigen accumulates on follicular dendritic cells (FDCs), key stromal cells in B cell follicles that support the GC response. These changes in cell- and tissue-level pharmacokinetics correlate with robust and sustained GC responses, greatly augmented affinity maturation, and enhanced development of plasma cells in the bone marrow. Thus, ALD vaccines appear promising for engineering multiple elements of vaccine kinetics that augment humoral immunity.

## RESULTS

### Atomic layer deposition preserves the structural integrity of complex HIV Env trimer immunogens

The process for encapsulation of vaccines in ALD microparticles is schematically shown in **Fig. 1A**: A liquid formulation of vaccine in buffer containing a glass-forming polysaccharide mixture is spray-dried to embed the vaccine within glassy microparticles, which are subsequently coated in a fluidized bed reactor with nanoscopic alumina layers by the atomic layer deposition process(*44*). Each cycle of the ALD process comprises two sequential, self-limiting reactions. In the first, trimethyl aluminum reacts with hydroxyl groups on particle surfaces, replacing the surface hydroxyl groups with methyl- and dimethyl aluminum oxides and liberating methane. Water vapor added to initiate the second reaction reacts with the methyl- and dimethyl-aluminum oxides to produce additional methane, leaving the surface coated with a single molecular layer of aluminum oxide with a thickness of ∼0.24 nm and regenerating the surface hydroxyl groups (*40*). Repeated cycles of this process build an alumina shell layer-by-layer up to a desired coating thickness.

**Fig. 1:**
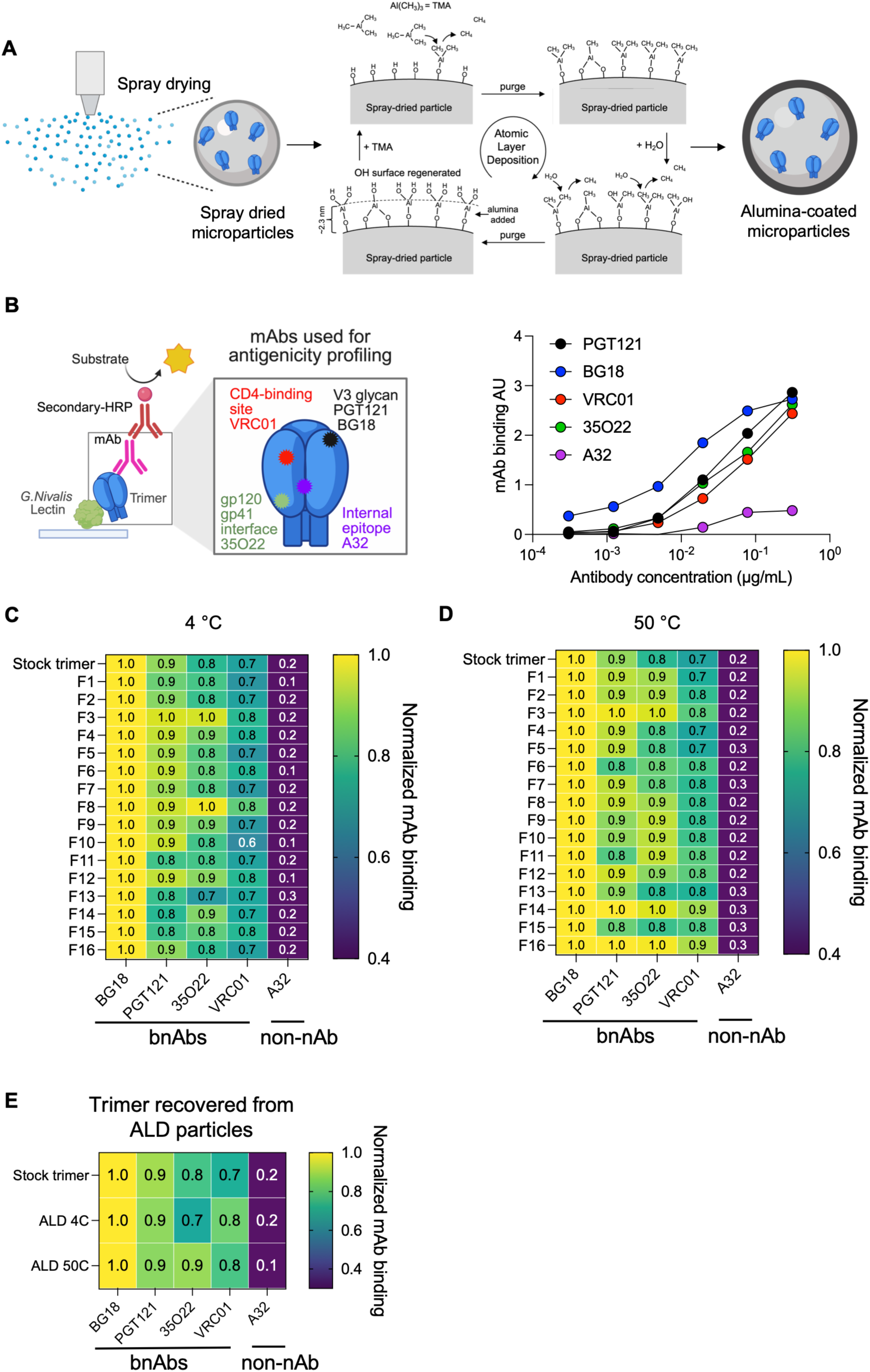
Stabilized HIV Env trimers are structurally intact after ALD particle processing. **A)** Schematic of ALD core-shell particle fabrication process. **B)** Schematic of Env trimer antigenicity profiling assay and representative raw ELISA binding data for stock protein antigen. **C-D)** Antigen was spray-dried in different excipient formulations (1–16) followed by subsequent incubation at 4 °C or 50 °C for 14 days. Samples were analyzed by the antigenicity profiling assay. Shown are area-under-the-curve values for the ELISA data normalized to values for BG18 binding. **E)** Antigen was spray-dried using formulation 1, coated with 50 layers of alumina, and ALD particles were incubated at 4 °C or 50 °C for 14 days. Subsequently, the alumina shell was dissolved, and integrity of encapsulated antigen was analyzed through antigenicity profiling as in c-d.

We previously showed that vaccination of mice with human papillomavirus 16 virus-like particles, rabies virus (RabiVax-S), or SARS-CoV-2 receptor-binding domain antigens encapsulated in ALD particles elicits neutralizing antibody responses comparable to or superior to bolus vaccinations, suggesting preservation of antigen integrity in ALD particles (*36, 38–40*), but direct analysis of the impact of ALD encapsulation on antigen structure has not been reported. Retention of the structural integrity of the antigen through the encapsulation process is critical, especially for HIV immunogens that are engineered to engage rare protective precursor B cells. Exploiting the availability of a broad range of structure-sensitive monoclonal antibodies that recognize distinct epitopes on HIV Env trimer immunogens, we evaluated the impact of spray drying and ALD coating on HIV trimer antigen structure. We employed the HIV immunogen N332-GT2 as a testbed antigen (*11*), as it is representative of a broad class of stabilized HIV Env trimers known as germline targeting immunogens, which are currently undergoing clinical development to activate bnAb-precursor B cells (*45*). We developed an antigenicity profiling assay, wherein N332-GT2 (hereafter, antigen for simplicity) is captured on lectin-coated ELISA plates and then incubated with different dilutions of a panel of monoclonal antibodies that recognize different structural epitopes on the antigen (**Fig. 1B**). bnAbs that bind to correctly-folded trimers show steadily increasing binding with increasing concentration, while a control antibody that recognizes an epitope on gp120 that is mostly occluded on properly folded trimers (A32) shows low binding (**Fig. 1B**). To quantify these binding characteristics, we calculated the area-under-the-curve of ELISA signal vs. antibody concentration and normalized the values to the binding by antibody BG18, a bnAb recognizing the V3 glycan site of vulnerability on HIV Env for which N332-GT2 was particularly engineered for high affinity recognition (*11*). This normalization also accounts for slight variations in the amount of antigen captured on the ELISA plates. To determine an optimal formulation for the initial spray-drying step, we used lyophilization to embed the antigen in a series of 16 different glassy trehalose formulations that varied in pH, ionic strength, and inclusion of secondary excipients (**Table S1**). To further provide insights into whether any of these formulations provided greater thermostability for the antigen in the dry, glassy state, each sample was subsequently incubated for 2 weeks either at 4°C or 50°C, then reconstituted in water and analyzed for structural integrity of the recovered antigen using the antigenicity profiling ELISA. As shown in **Fig. 1C-D**, most of the tested formulations showed relatively well-preserved structural integrity of the antigen, with high levels of binding of each of the tested bnAbs and low binding of A32. We selected formulation 1 for further studies based on its favorable antigenicity profile at both test temperatures, employing trehalose as the glassy matrix (**Table S1**), and fixed the antigen concentration at 0.08 wt% antigen per mg spray-dried particles.

We next carried out spray-drying followed by atomic layer deposition on this selected formulation, coating particles with 50-300 layers of alumina (hereafter, we indicate the number of alumina layers by a number following the atomic layer deposition abbreviation, e.g., ALD50 = 50 layers of alumina). Consistent with prior studies, the resulting core-shell particles were polydisperse, with number-average diameters, D_10_, and D_90_ values of 5.1 µm, 2.3 µm, and 8.5 µm, respectively (**fig. S1A-B**). Focused ion beam scanning electron microscope (FIB-SEM) analyses of particles coated with 50, 100, or 200 layers of alumina showed thicknesses corresponding to 2.4±0.2 Å/ALD cycle (**fig. S1C-D**). Thus, even at the highest number of alumina coats applied here (300), the alumina shell thickness is very thin compared to the diameter of spray-dried particle core, thus minimizing any potential particle size effects on shell dissolution rates due to surface curvature. We incubated coated particles at 4°C or 50°C for 2 weeks, then dissolved the alumina coating in citrate/phosphate buffer and carried out antigenicity profiling on the recovered antigen. Antigen recovered from ALD50 particles showed well preserved structural integrity similar to the spray-dried product and unmanipulated stock protein at both test temperatures (**Fig. 1E**). Altogether, these data indicate that ALD microparticles can encapsulate and release these engineered immunogens with a high degree of structural fidelity.

### Interactions of ALD particles with antigen presenting cells *in vitro*

A portion of ALD particles are of a size that would be compatible with phagocytosis by antigen presenting cells (APCs) such as macrophages and dendritic cells, and thus we first examined interactions of ALD particles with these APCs *in vitro*. RAW 264.7 macrophages were incubated with ALD30 particles encapsulating antigen labeled with an Alexa Fluor 647 dye. This very thin ALD coating is expected to begin releasing its contents almost immediately and provided the opportunity to observe potential digestion of particles in phagosomes. After 24 hr of incubation, RAW cells exposed to suspensions of ALD30 were positive for particle uptake (**Fig. 2A**). While many cells could be observed that appeared to have internalized intact particles, we also observed cells positive for antigen signal without a clear microparticle present, and cells that appeared to have internalized particle shells that retained a residual rim of trimer signal but that had released their contents (**Fig. 2A-B**). We repeated this experiment using ALD50 particles loaded with the small-molecule dye calcein, and quantified uptake as a function of particle concentration by flow cytometry. The fraction of the macrophage population positive for particle uptake appeared to saturate at approximately 90% at particle concentrations of 700 μg/mL or higher (**Fig. 2C**).

**Fig. 2:**
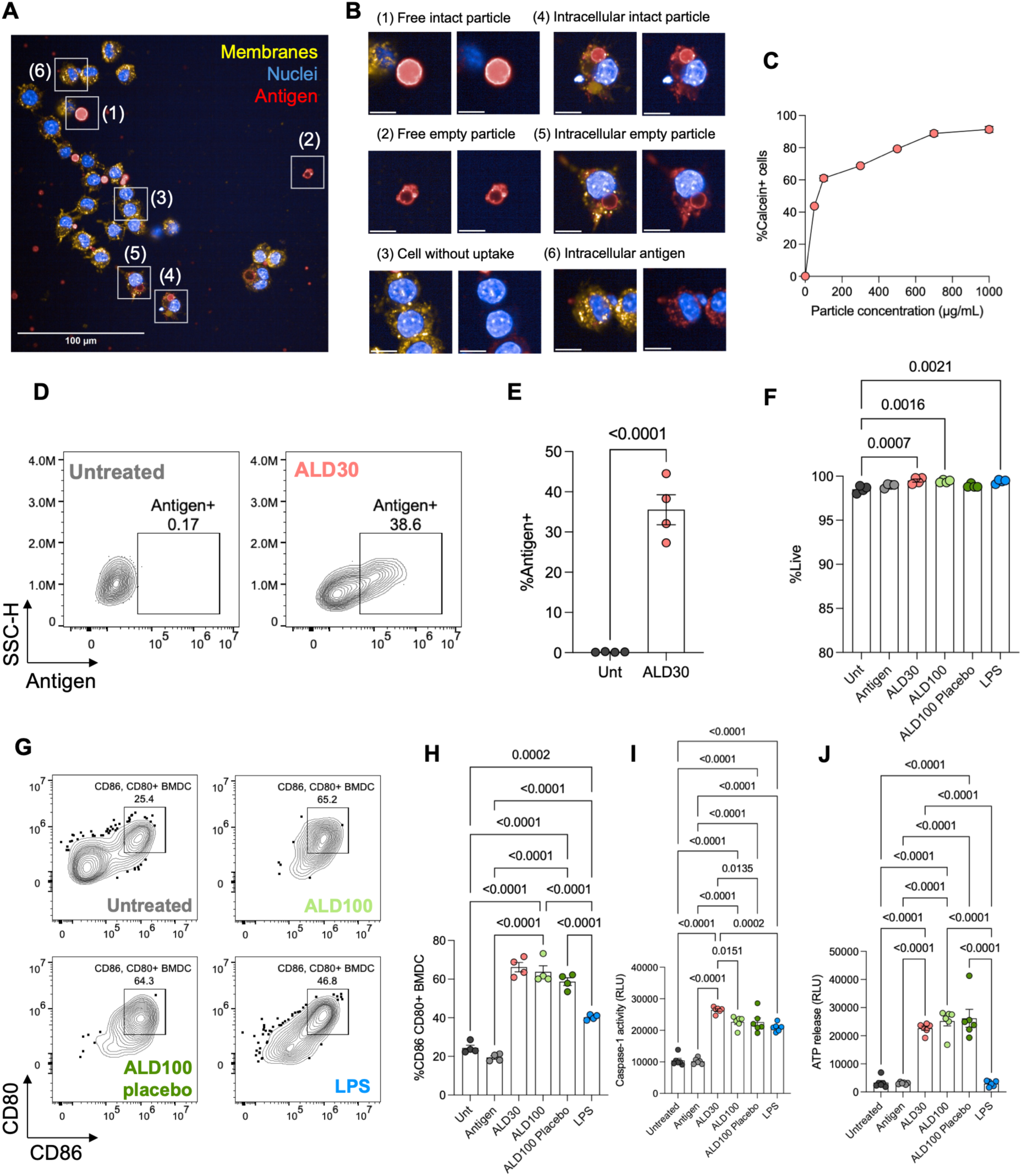
ALD particles are taken up by phagocytes and activate dendritic cells *in vitro*. **A-B)** Activated RAW 264.7 cells were incubated for 24 hr with 10 μg/mL suspensions of ALD30 microparticles containing Alexa Fluor 647-labeled antigen and imaged to assess particle internalization. Shown is a representative micrograph **(A)** and magnified views of several cell-particle states **(B)**. **C)** Activated RAW 264.7 cells (*n* = 3 samples) were incubated for 2 hr with indicated concentrations of calcein-containing ALD50 microparticles and analyzed by flow cytometry for quantification of mean calcein^+^ cells. **D-H)** BMDCs (*n* = 4 samples/group) were left untreated or incubated with 5 µg/mL free Alexa Fluor 647-labeled antigen, 0.5 mg/mL ALD30 or ALD100 particles encapsulating labeled antigen, 0.5 mg/mL empty (placebo) ALD100 particles, or 10 ng/mL LPS for 24 hr, followed by flow cytometry analysis. Shown are representative flow cytometry plots of antigen uptake **(D)**, quantification of antigen^+^ cells **(E)**, viability of cells **(F)**, representative flow cytometry plots of CD80/CD86 expression on BMDCs **(G)** and corresponding quantification **(H)**. **I-J)** BMDCs (*n* = 6 samples/group) were incubated with antigen-loaded ALD30 particles or LPS as in **(D)**, and caspase-1 activity **(I)** or ATP release **(J)** were assessed after 24 hr. Shown are means ± s.e.m. Significance was determined using two-tailed unpaired t-test **(E)** and one-way ANOVA with Tukey’s post hoc analysis **(F, H-J)**. Scale bars correspond to 10 μm **(B)**.

We next analyzed the interaction of ALD particles carrying fluorescently-labeled antigen with primary mouse bone marrow-derived dendritic cells (BMDCs) *in vitro* and compared stimulation of BMDCs with the potent Toll like receptor-4 agonist lipopolysaccharide (LPS) as a positive control. Incubation of BMDCs for 24 hr with ALD30 particles encapsulating Alexa Fluor 647 labeled antigen led to particle uptake in ∼35% of the cells as determined by flow cytometry (**Fig. 2D-E**). In contrast, soluble antigen added to the cells at an equivalent antigen concentration as ALD particles showed minimal uptake (**fig. S1E**). We then analyzed effects of ALD particle uptake on BMDCs, comparing ALD30, ALD100, and “empty” ALD100 particles lacking encapsulated antigen. Particle uptake was non-toxic (**Fig. 2F**), but DCs treated with ALD particles (but not free antigen) were activated as evidenced by upregulation of the costimulatory receptors CD80 and CD86 to a level comparable to treatment with LPS (**Fig. 2G-H**). Microparticles of diverse materials are known to activate inflammasomes in phagocytes (*28, 46, 47*), and thus we analyzed the activation of caspase-1, a key player in signaling downstream of the NLRP3 inflammasome (*46*). Both ALD particles and LPS treatment induced caspase-1 signaling in BMDCs (**Fig. 2I**). Note that caspase activation can occur in DCs without necessarily leading to cell death, through a process known as DC hyperactivation (*48–50*). However, caspase-1 activation promotes the release of ATP from inflammasome-triggered cells. Measurement of ATP in culture supernatants revealed that ALD particles (but not LPS) induced robust ATP release from treated cells (**Fig. 2J**), which itself can strongly activate DCs through purinergic receptors (*51, 52*). These responses were not different for ALD particles with different alumina shell thickness and did not depend on the presence of the encapsulated antigen. Thus, the alumina surfaces of ALD particles activate elements of the inflammasome pathway known to contribute to DC activation and promotion of vaccine immunity.

### Alumina shell thickness programs antigen pharmacokinetics and initiation of the *in vivo* immune response

The premise of ALD core-shell particle design is that vaccine release kinetics post-injection are dictated by the alumina shell, which blocks dissolution of the polysaccharide/antigen core until the shell has dissolved (**Fig. 3A**). However, the *in vivo* pharmacokinetics of antigens administered in ALD-coated particles have remained unclear. To gain insight into the impact of ALD delivery on antigen persistence at the injection site, we injected suspensions of fluorophore-labeled antigen encapsulated in ALD particles subcutaneously near the tail base of albino C57BL/6 mice and imaged the clearance of antigen over time by *in vivo* whole-animal fluorescence imaging. To slow particle settling and ease administration, we resuspended ALD particles in a biodegradable triglyceride oil for injection, but we found no difference in antigen clearance kinetics whether the particles were injected in tris buffer or suspended in oil (**fig. S1F**), and the oil rapidly cleared from the injection sites within 7 days. As ALD vaccines have previously been shown to be substantially more potent than immunization with traditional alum (*40, 41*), we benchmarked ALD delivery against soluble antigen mixed with a saponin adjuvant (SMNP), an established highly potent bolus immunization comparator for this immunogen that is currently being evaluated with HIV vaccines in the clinic (*19, 21, 22, 53, 54*). As previously observed (*55, 56*), immunization with SMNP-adjuvanted antigen led to rapid clearance from the injection site (**Fig. 3B-C**). In contrast, antigen delivered in ALD particles exhibited multi-phasic clearance kinetics: Initially, delayed antigen clearance was observed as evidenced by a period of minimal signal decay, and duration of this delay was dictated by alumina shell thickness (**Fig. 3B-C**). Following the initial delay, a period of more rapid, steady clearance of labeled antigen from the injection site was observed that continued for 30-50 days (**Fig. 3B-C**). This was followed in each case by a final plateau of slow antigen clearance, which we hypothesize reflects residual antigen released from particles but trapped at the injection site. To estimate the delay time before sustained clearance began for each sample, we fit the initial plateau region and sustained clearance regions of the curves via linear regression and calculated the intersection of these two lines (**fig. S1G**). This analysis showed a direct linear correlation between the thickness of the alumina shell and the delay until the beginning of sustained clearance, demonstrating control of the timing of vaccine release *in vivo* by the ALD coating thickness (**Fig. 3D**). To correlate this pharmacokinetics to immune response, we immunized mice with ALD particles loaded with unlabeled antigen and tracked serum antibody responses over time. Antigen-specific IgG titers following bolus vaccination with soluble antigen reached values of ∼10^3^ by 7 days, while ALD50 and ALD100 injections took between ∼2-3 weeks to reach these values, and further increased over the subsequent 30 days (**Fig. 3E**). IgG titers following administration of ALD300 particles were low until after day 40 when the sustained clearance phase began, and reached titers of 10^4^ around day 100 (**Fig. 3E**). Thus, ALD particles can be used to both program a delay in the release of vaccine antigen and greatly prolong antigen exposure once release begins relative to traditional bolus vaccination.

**Fig. 3:**
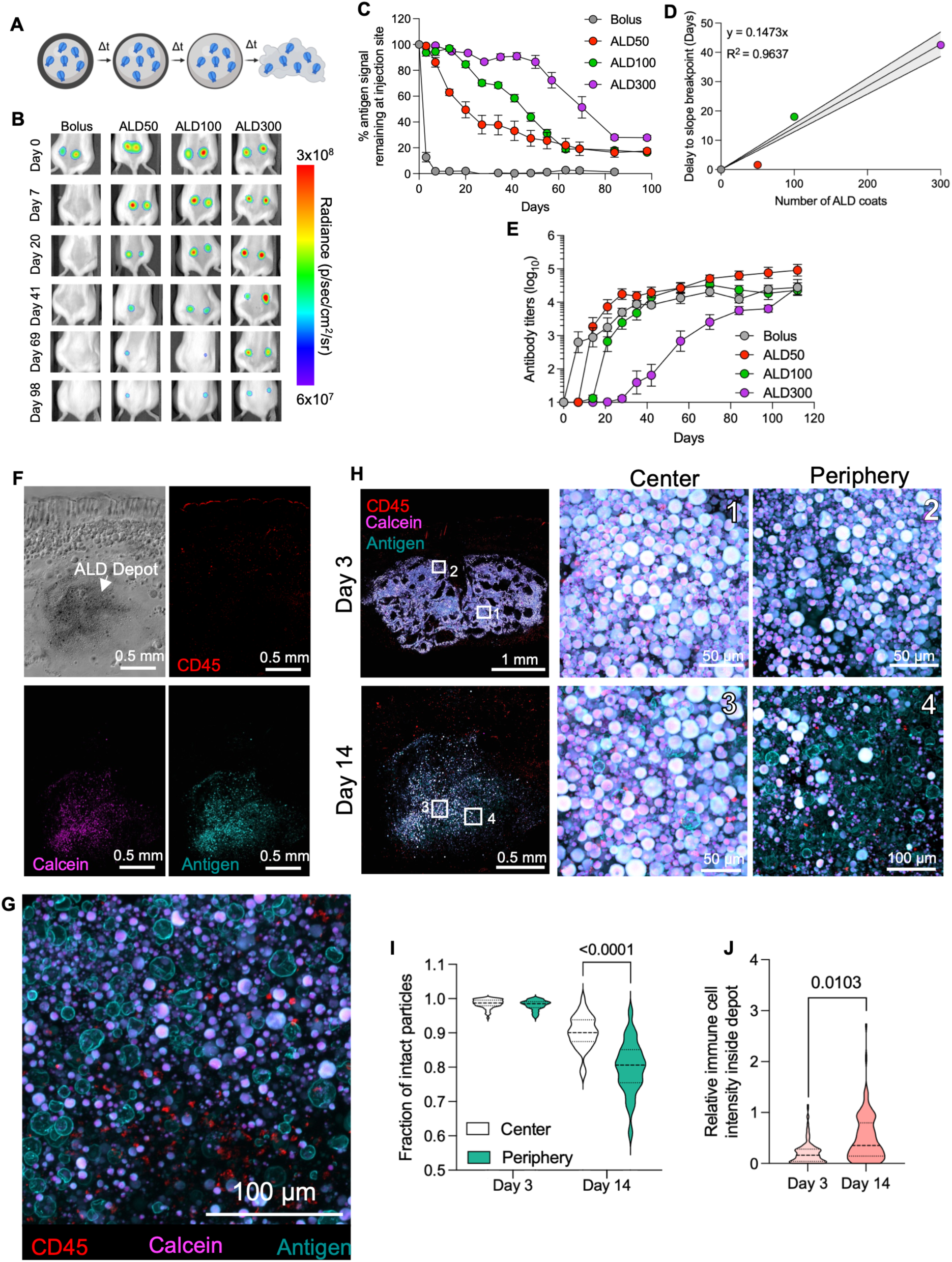
ALD particles show multi-phase antigen release programmed by the alumina shell thickness. **A)** Schematic illustrating dissolution of alumina shell and subsequent antigen release from ALD core-shell microparticles. **B-D)** Albino C57BL/6 mice (*n* = 5 animals/group) were injected s.c. with 10 µg Alexa Fluor 647–labeled antigen administered either as a liquid bolus with 5 μg SMNP saponin adjuvant or encapsulated within ALD particles with varying alumina shell thickness. ALD particles were suspended in triglyceride oil for injection. Antigen persistence at the injection site was monitored longitudinally by whole-animal fluorescence imaging. Shown are representative photograph/fluorescence signal overlays **(B)**, total fluorescence signal over at the injection site over time **(C)**, and relationship between the number of alumina layers deposited on particles by ALD and the time when rapid antigen clearance initiates **(D)**. **E)** C57BL/6 mice (*n* = 5 animals/group) were immunized with 5 µg antigen administered as a liquid bolus with 2.5 μg SMNP adjuvant or as ALD particles suspended in oil and serum IgG titers were tracked over time. **F-J)** C57BL/6 mice (*n* = 4 animals/group) were immunized with 5 µg Alexa Fluor 647-labeled antigen and 7.5 μg calcein administered as ALD particles suspended in oil. Injection sites were excised and visualized using fluorescence microscopy. Shown are representative low-magnification images of the injection site as brightfield and single-channel fluorescence for CD45 staining (red), calcein (magenta), and antigen (cyan) at day 14 **(F),** and high magnification images of the periphery at day 14 **(G)**. Also shown are representative images of ALD particles at the depot periphery and center at days 3 and 14 post injection **(H)**, fraction of intact particles at the periphery and center **(I)** and immune cell infiltration inside the depot **(J)**. Error bars represent s.e.m. Statistical significance was assessed using nested one-way ANOVA with Šidák’s multiple-comparison test **(I)** and nested two-tailed t-test **(J)**.

Two questions arising from the IVIS data were (1) is antigen primarily released in the extracellular space or within immune cells recruited to the injection site? and (2) what governs the duration of the rapid/steady clearance phase once ALD particles begin releasing antigen? To gain further insight, we examined the fate of ALD particles at the injection site by immunofluorescence microscopy. C57BL/6 Mice were immunized with ALD50 particles co-loaded with Alexa Fluor 647-labeled antigen and the small molecule fluorophore calcein. We reasoned that particles positive for both fluorescence signals would definitively represent ALD shells that had not opened to release their contents, because the calcein dye would be expected to rapidly clear from the injection site on release. Imaging of thick microtome histological sections collected from injection sites two weeks post immunization revealed an obvious depot of ALD particles, many of which were still unopened, consistent with the IVIS data (**Fig. 3F**). Higher resolution imaging of thin cryostat tissue sections near the edge of the depot strikingly revealed collections of ALD particles that had opened and released their contents, which were marked only by a thin “ghost” of the particle shell that had adsorbed some fluorescent antigen (**Fig. 3G** and **fig. S1H**). A modest infiltration of CD45^+^ immune cells was also observed, though a majority of the intact ALD particles appeared to be extracellular (**Fig. 3G** and **fig. S1H**). Imaging at two time points revealed distinct structural changes across the depot. At day 3, prior to substantial ALD50 release, particles appeared intact throughout the center and periphery of the depot confirmed by colocalization of calcein and Alexa Fluor 647 signal (**Fig. 3H-I** and **fig. S1I**). In contrast, by day 14, when approximately 40% of the antigen had cleared from the injection site (**Fig. 3C**), the periphery of the particle depot contained significantly more empty shells than the center (**Fig. 3H-I**). Further, quantification of CD45 staining intensity within the depot relative to surrounding tissue indicated increased immune cell infiltration by day 14 (**Fig. 3J** and **fig . S1J**). Altogether, these observations suggest that a substantial proportion of antigen release from ALD particles occurs extracellularly. We observed only fully “loaded” particles and completely empty shells, suggesting that *in vivo*, individual particles release over a narrow time window. However, the distribution of opened capsules suggests the very low solubility of aluminum in physiologic conditions limits dissolution/opening of particles near the center of injected ALD vaccine depots until shells at the periphery have dissolved and cleared, thus widening the time window over which vaccine release occurs once particles begin to open.

### ALD-delivered antigen progressively accumulates in APCs at the injection site and in draining lymph nodes

We next examined the response of innate immune cells to ALD or soluble antigen vaccines at the injection site and draining lymph nodes over time. Groups of mice were immunized with either soluble dye-labeled antigen mixed with SMNP or ALD50 particles carrying the same antigen and no additional adjuvant, and cells present at the injection site were analyzed by flow cytometry at different time points post-immunization (**Fig. 4A, fig. S2A**). One-week post-immunization, the most prominent cell types recruited to the injection site were monocytes and macrophages for both soluble and ALD vaccines, with ALD also eliciting a substantial accumulation of neutrophils (**Fig. 4B**). Examining the response of these cells over time, bolus immunization led to a peak in the number of monocytes and neutrophils recruited to the injection site 2 days post immunization, followed by a decline to the pre-immunization baseline by day 14 (**Fig. 4C-E**). By contrast, monocyte and neutrophil levels observed at the injection site following ALD vaccination continued to increase through day 7 and were still elevated at day 14 (**Fig. 4C-E**). More strikingly, bolus immunization led to a transient population of antigen-positive immune cells that could be detected 2 days post immunization and decayed to baseline by day 7, whereas ALD50 vaccination led to antigen-positive immune cell populations observed at day 2 that were similar to those produced by bolus injection but that remained elevated or even increased through day 14 (**Fig. 4F-H**). Although recruited at lower absolute numbers, similar trends of increasing antigen acquisition over time were also detected in cDC1 dendritic cells, cDC2 dendritic cells, and monocyte-derived dendritic cells (moDCs) following ALD vaccination (**fig. S2B-D**). A proportion of these antigen presenting cells taking up antigen at the injection site showed early signs of activation, as indicated by upregulation of the costimulatory receptor CD40 (**fig. S2E**).

**Fig. 4:**
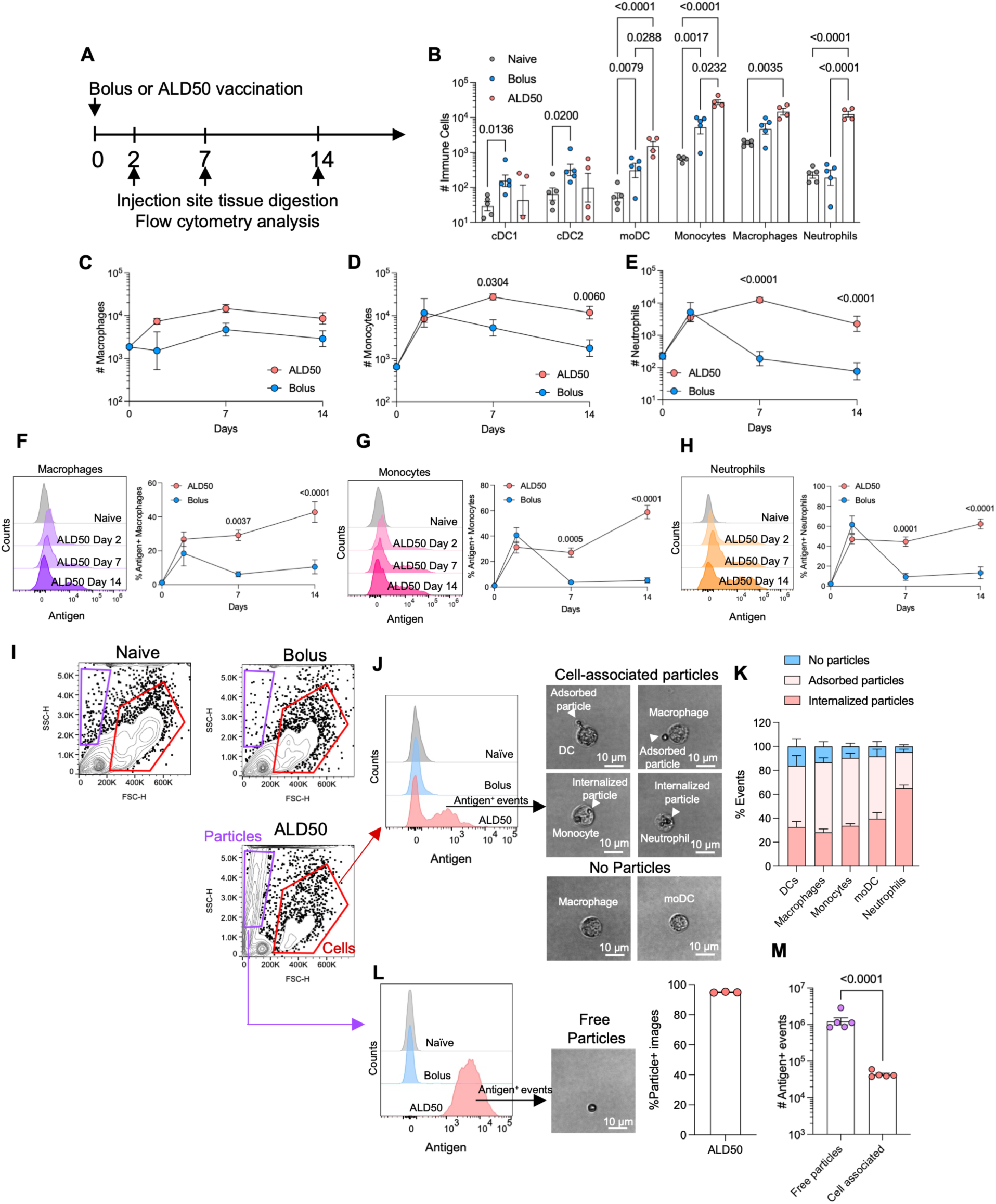
ALD promotes immune cell infiltration and antigen uptake at injection site. C57BL/6 mice (*n* = 5 animals/group) were immunized s.c. with 10 µg AlexaFluor 647–labeled antigen delivered either as a liquid bolus with 5 μg SMNP adjuvant in PBS or encapsulated within ALD50 particles and resuspended in triglyceride oil. Skin and s.c. tissue at the injection site was digested for flow cytometry analysis of recruited immune cells at different time points post immunization. **A)** Timeline of experiment. **B)** Enumeration of innate immune cells at day 7. **C-E)** Enumeration of macrophages, monocytes, and neutrophils at the injection site over time. **F-H)** Representative histograms of antigen uptake and frequencies of antigen^+^ cells over time for macrophages **(F)**, monocytes **(G)**, and neutrophils **(H)**. **I-M)** C57BL/6 mice (*n* = 5 animals/group) were immunized with 10 µg antigen as a liquid bolus with 5 μg SMNP adjuvant or as ALD50 particles suspended in oil. At day 14 post immunization, injection site tissues were isolated, digested into single-cell suspensions, and analyzed by imaging flow cytometry. Shown are representative forward/side scatter flow cytometry plots **(I)**, example gating for antigen^+^ live CD45^+^ cells with representative brightfield images of analyzed cells **(J)**, summary analysis of antigen^+^ cells (compiled from over 1,000 total flow cytometer images) **(K)**, representative histograms of antigen signal in “free particle” FSC/SSC gate alongside example event image and enumeration of particle^+^ images **(L)**, and summary enumeration of antigen^+^ events in the free particle vs. cell-associated gates **(M)**. Error bars represent s.e.m. Statistical significance was assessed using two-way ANOVA with Tukey’s **(B)** and Šidák’s **(C-H)** multiple comparison test and unpaired two-tailed t-test **(M)**.

To gain further insights into the interactions of immune cells infiltrating at the injection site and ALD particles, we injected fluorescently labeled antigen either encapsulated in ALD50 particles or administered as a bolus vaccine and then harvested and digested the injection site for flow cytometry analysis 2 weeks later, using a flow cytometer equipped with a camera for brightfield imaging of each analyzed cell/event. We defined a broad mid-to-high FSC/SSC flow cytometry gate to capture all cells (**Fig. 4I**). Further subgating on live, CD45^+^ immune cells (**fig. S2F**), a significant proportion of immune cells at the ALD50 injection site were positive for antigen, as discussed above (**Fig. 4J**). To determine whether these cells had taken up antigen still encapsulated in ALD particles vs. free antigen, we examined brightfield images of the live CD45^+^antigen^+^ events, further phenotyping various myeloid cell populations by gating on cell surface markers. Event imaging revealed 3 populations: cells that appeared to have internalized one or more particles, cells with particles adsorbed to their surfaces, and cells that had no apparent particle association (**Fig. 4J**). We observed each of these 3 classes of events in all major myeloid cell populations including macrophages, monocytes, neutrophils, and dendritic cells (**Fig. 4J** and **fig . S2G**). Enumeration of cells in each category revealed that the majority (>80%) were associated with intact particles either internalized or adsorbed, with neutrophils showing slightly higher internalized particle proportions compared to the other myeloid cells (**Fig. 4K**).

Flow cytometric analysis of enzymatically-digested injection site tissue from ALD-injected animals also showed a substantial population of events at low forward scattering intensity (FSC)/high side scattering intensity (SSC), which were brightly positive for antigen; events in this gate were antigen-negative in naïve or bolus-immunized mice, leading us to hypothesize that these might be cell-free particles (**Fig. 4I, L**). Imaging cytometry confirmed that these low FSC/high SSC events in the ALD-immunized animals were indeed free particles (**Fig. 4L**). Despite the fact that the total number of free particle events detected in the cytometer represents a minor fraction of the total particle number injected (likely reflecting a combination of particle dissolution during incubation at the injection site and settling/loss of particles in the tissue digestion process), free particles greatly outnumbered particles taken up by cells (**Fig. 4M**), consistent with the qualitative observations made via histology.

In parallel to analysis of the injection site, we characterized the distribution of antigen and activation of APC populations in the draining lymph node (**fig. S3A**). At 2 days post immunization with bolus antigen and SMNP adjuvant (a timepoint near the peak of both lymph node activation and antigen accumulation for soluble antigen/adjuvant immunization (*53, 57*)), antigen could only be detected clearly above background in ∼10% of migratory cDC2 cells (e.g., DCs that have migrated from the injection site, (**Fig. 5A**); antigen levels then decayed at later time points (not shown). Strikingly, at day 21 for ALD50 immunization, a time point when ∼50% of ALD50-administered antigen has been cleared from the injection site, 30% of monocyte-derived DCs (moDCs), 40% of lymph node-resident cDC2, and 60% of cDC1 were antigen^+^ (**Fig. 5A**). Histology of draining lymph nodes at day 14 post immunization with ALD50 particles carrying fluorescent antigen revealed a substantial accumulation of antigen^+^ cells in the medullary sinuses at the center of lymph nodes, with scattered antigen^+^ cells in the T cell areas and near B cell follicles (**Fig. 5B**). To assess whether these cells contained intact particles at this time point, we carried out imaging cytometry on dLNs, and found that distinct from the injection site, only 5-10% of antigen^+^ APCs showed evidence for intact particles at this timepoint (**Fig. 5C-D, fig. S3B**). This could reflect that antigen released extracellularly at the injection site was trafficked through lymph to these resident APCs in the lymph node, or that particles had been trafficked to the node and digested intracellularly by this time.

**Fig. 5:**
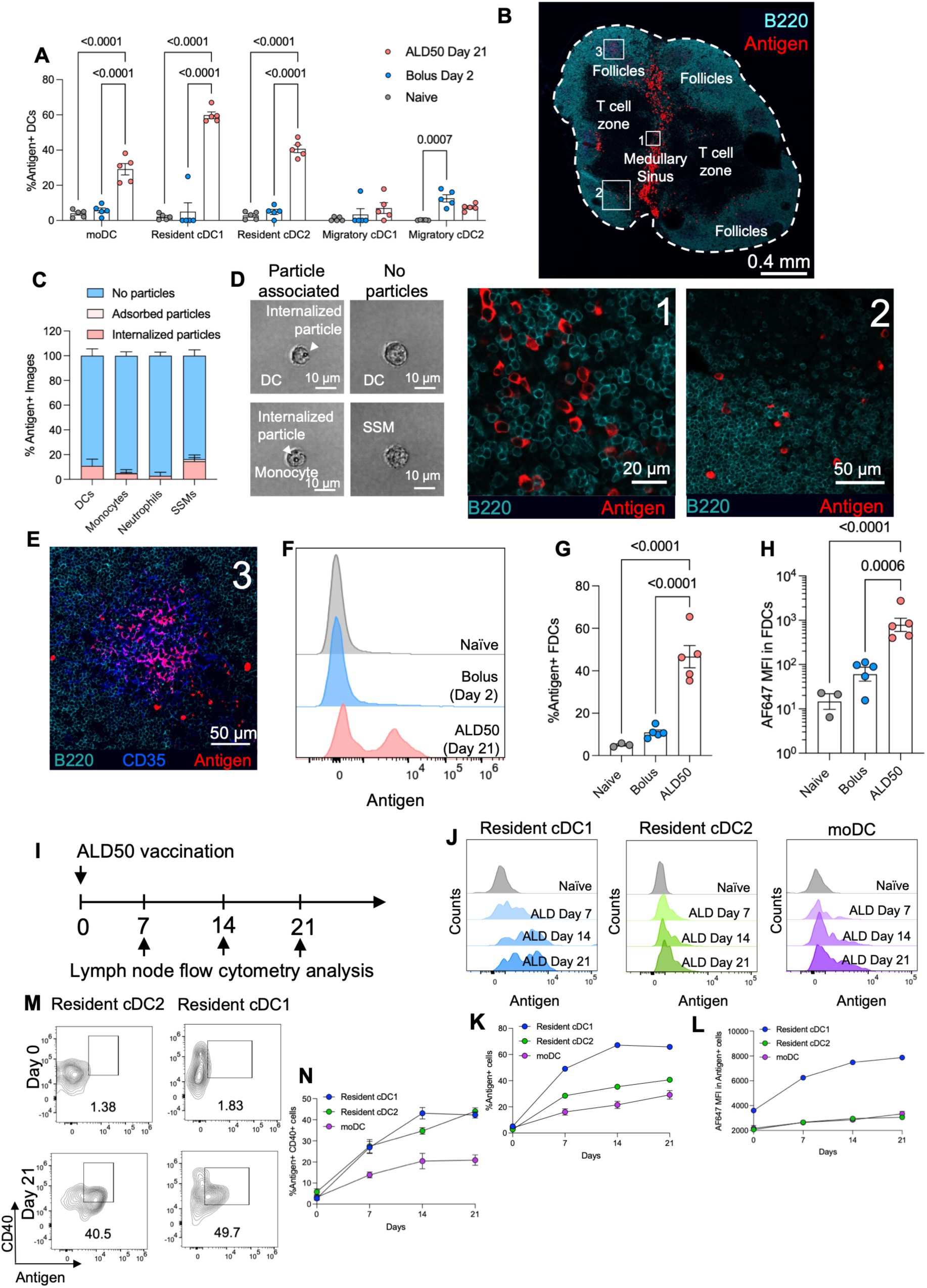
ALD particles promote antigen uptake in lymph node DC populations and deposit antigen on follicular dendritic cells. C57BL/6 mice (*n* = 5 animals/group) were s.c. immunized with 10 μg Alexa Fluor 647–labeled antigen delivered either as a liquid bolus dose with 5 μg SMNP adjuvant in PBS or encapsulated within ALD50 particles with 50 coats of alumina and resuspended in triglyceride oil. Lymph nodes were harvested for flow cytometry or histology analysis. **A)** Frequencies of antigen^+^ events in key dendritic cell subsets. **B)** Representative histology of draining lymph nodes at day 14 post ALD50 immunization. **C-D)** Imaging cytometry analysis of antigen^+^ cells from dLN at day 14. **E)** Histological imaging of FDC network inside a B cell follicle of the LN. **F-H)** Antigen accumulation on FDCs. Shown are representative histograms of antigen uptake on FDCs **(F)**, percent antigen^+^ FDCs **(G)**, and mean fluorescence intensity (MFI) of antigen signal on FDCs **(H)** at day 2 post bolus or day 21 post ALD50 immunization. **I-M)** Distribution of antigen among key DC populations. Shown are timeline of tissue collections **(I)**, representative histograms of antigen accumulation over time in LN-resident DCs and moDCs **(J)**, mean antigen^+^ DC frequencies over time (**K**), antigen mean fluorescent intensities (MFIs) in DCs over time **(L)**, representative flow cytometry plots of expression of activation marker CD40 in antigen^+^ DCs **(M)**, and mean frequencies of antigen^+^ activated DCs over time **(N)**. Error bars represent s.e.m. Statistical significance was assessed using two-way ANOVA with Tukey’s post-hoc test **(A)**, and one-way ANOVA with Tukey’s post-hoc test **(G, H)**.

Continuous delivery of antigen during the first few weeks of an immunization has been shown to lead to capture of antigen on follicular dendritic cells (FDCs) (*20, 58–60*), key stromal cells that reside in B cell follicles and present antigen over time to B cells during an ongoing germinal center (GC) response. Histology analysis revealed substantial antigen accumulation on FDCs in B cell follicles (**Fig. 5B, E**). Flow cytometry analysis of FDCs isolated from pooled lymph nodes (**fig. S3C**) at day 2, representing the time for peak antigen accumulation following bolus antigen immunization, showed no detectable deposition of antigen on these cells over the naïve control **(Fig. 5F-H**). By contrast, lymph nodes harvested at day 21 following ALD50 vaccination showed substantial antigen capture on FDCs (**Fig. 5F-H**) (*60*). Thus, ALD vaccination led to substantial accumulation of antigen in APCs in the medullary sinuses and in FDCs within B cell follicles, as well as uptake in cells scattered through the T cell zones.

We next characterized the kinetics of antigen accumulation in LN APCs following ALD50 immunization (**Fig. 5I**). Each DC population showed increases over 3 weeks in both the fraction of antigen^+^ cells and the amount of antigen acquired per cell, with 40% of resident cDC2 and 60% of resident cDC1 acquiring antigen by day 21 (**Fig. 5J-L**). Further, examination of costimulatory receptors showed that a portion of these antigen^+^ cells were strongly activated (**Fig. 5M-N**). Altogether, ALD delivery greatly increased the accumulation and persistence of antigen in key APC populations over time compared to traditional bolus antigen/adjuvant immunization.

Given the local immune activation observed in the injection site and dLNs, we also evaluated whether ALD particles cause systemic cytokine elevation. Across multiple timepoints post-vaccination, ALD particles did not induce systemic upregulation of any of the cytokines measured, suggesting they do not trigger an acute systemic inflammatory response (**fig. S3D**). In contrast, SMNP administration produced moderate upregulation of IL-6 and CXCL10 at day 2 post-injection, as previously reported (*53*). IFN-γ, CXCL1, TNF-α, CCL2, IL-12p70, CCL5, IL-1β, GM-CSF, IL-10, IFN-β, and IFN-α were always below detection limits for both treatment groups.

### ALD vaccination promotes long-lived GCs and potent immune responses

Efficient recruitment and expansion of antigen-specific B cells into germinal centers (GCs) is thought to be critical for priming bnAb responses to HIV (*61, 62*), as GCs are the site where B cells accumulate mutations that permit high affinity antigen binding, and are also the source of high-affinity long-lived plasma cells that maintain protective antibody levels over time. We thus sought to assess the impact of sustained antigen accumulation in dLNs induced by ALD vaccines on GC responses to Env immunization. Mice were administered either bolus vaccine with antigen and SMNP, or antigen delivered in ALD50 particles, and cohorts were sacrificed at serial time points to track GC responses over 8 weeks post immunization (**fig. S4**). Bolus antigen immunization initiated a GC response that was sustained for ∼4 weeks and then declined to near baseline by 8 weeks (**Fig. 6A**). ALD50 vaccination by contrast led to a GC response that steadily increased over 4 weeks, consistent with the timeframe of antigen release; this GC response was still ongoing at 8 weeks (**Fig. 6A**). Interestingly, despite the distinct trajectories of GC responses, follicular helper T cells that help drive the GC response were primed with similar kinetics for the 2 vaccinations, with an early peak expansion at 1 week that then contracted thereafter (**Fig. 6B**). Short-lived plasmablasts that produce transient early antibody responses peaked sharply at 1 week for bolus immunization, while appearing later following ALD50 vaccination (**Fig. 6C**).

**Fig. 6:**
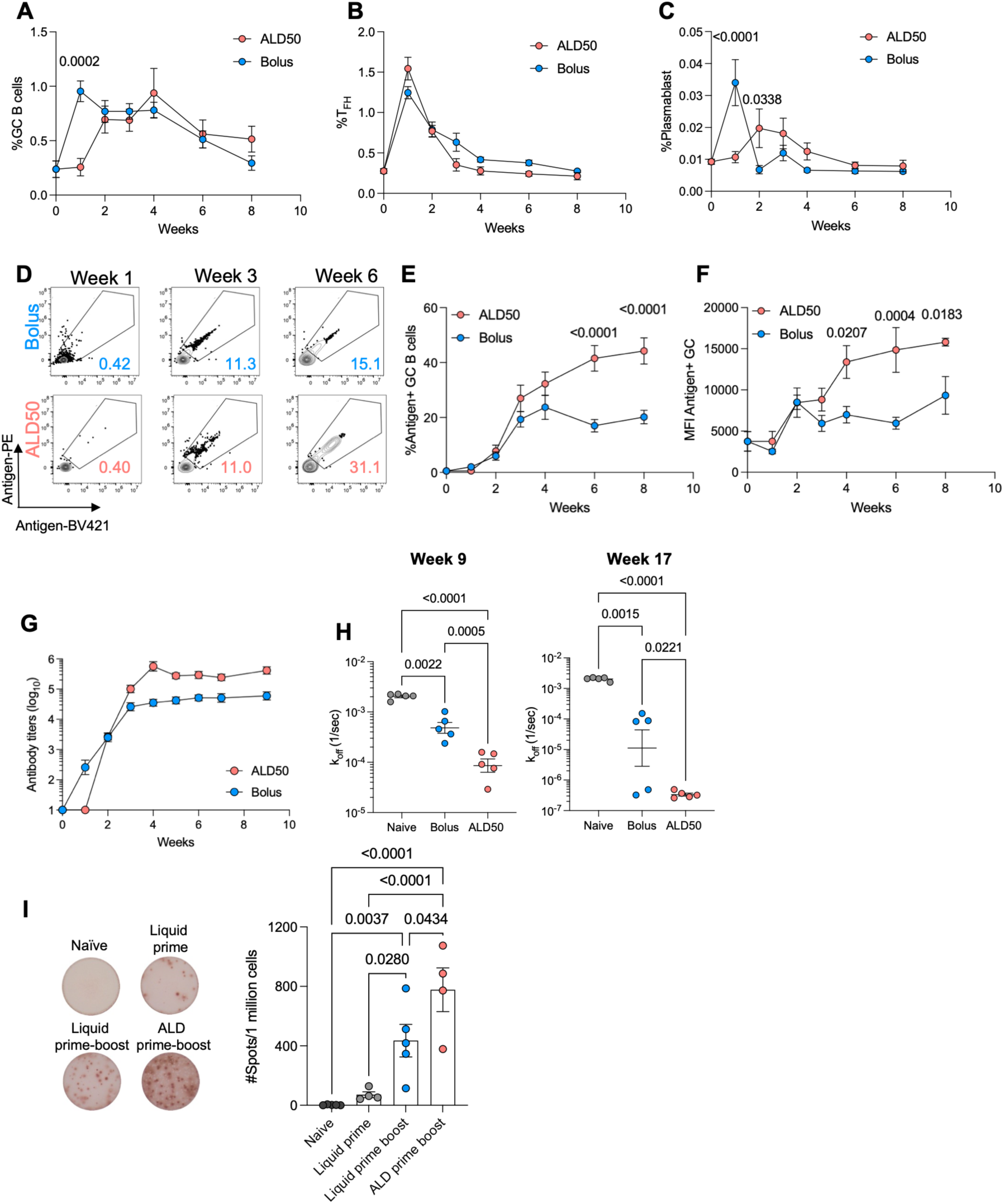
ALD delivery promotes long-lived germinal centers and enhanced affinity maturation of the antibody response. **A-G)** C57BL/6 mice (*n* = 5 animals/group) were s.c. immunized with 10 µg antigen delivered either as a liquid bolus dose with 5 μg SMNP adjuvant or encapsulated within ALD50 particles suspended in triglyceride oil. LNs were harvested at selected time points over time for flow cytometry analysis. Shown are percentages of B cells in GCs **(A)**, percentage of follicular helper cells among CD4^+^ T cells **(B)**, percentage of plasmablasts among B cells **(C)**, representative flow plots of fluorescent antigen tetramer staining among GC B cells **(D)**, percentage of antigen^+^ GC B cells **(E)**, and mean fluorescence intensity (MFI) of antigen tetramer staining on antigen^+^ GC B cells **(F)** over time. **G-H)** C57BL/6 mice (*n*=5 animals/group) were immunized as in (a) and serum IgG titers were tracked over time polyclonal IgG by ELISA **(G)** and serum was isolated at 9 or 17 weeks post immunization for biolayer interferometry analysis of antigen-specific antibody binding off rates **(H)**. **I)** To assess immune responses after a booster dose, C57BL/6 mice were immunized s.c. with 5 μg antigen and 2.5 μg SMNP on day 0; one cohort received a dose-matched booster at week 3. Additional groups were immunized with a mixed and equal weight amount of ALD30/ALD100 formulation on day 0. Antigen-specific antibody-secreting cell responses in the bone marrow measured at week 16 by ELISPOT. Error bars represent s.e.m. Statistical significance was determined by two-way ANOVA with Šidák’s multiple-comparison test **(A-C, E-F)** and one-way ANOVA with Tukey’s post-hoc test **(H, I)**.

We next examined the evolution of antigen-specific cells in the GCs. The frequency of antigen-binding GC B cells increased in the bolus group through week 3 but then plateaued at ∼20% of GC B cells (**Fig. 6D-E**). By contrast, antigen-binding GC cells in the ALD group continued to increase in frequency throughout the entire time course, reaching ∼2-fold higher levels by week 8 compared to the bolus group (**Fig. 6D-E**). In parallel, the brightness of antigen staining, a proxy for BCR affinity for the antigen, was higher in the ALD group from week 2 onwards (**Fig. 6D, F**).

To determine if these alterations in germinal centers elicited by bolus vs. ALD vaccination were also reflected in the downstream antibody response, we analyzed polyclonal antibodies induced by vaccination with 10 µg or antigen as a bolus or ALD50 vaccination (Note this is double the antigen dose used in the experiment shown in **Fig. 3E**). Serum antibody responses elicited by ALD50 lagged behind the bolus immunization but surpassed their titers between 2-3 weeks and plateaued about a log higher (**Fig. 6G**). We assessed the off-rate of binding to antigen using biolayer interferometry for polyclonal antigen-specific IgG isolated from sera of mice from each group at 9 or 17 weeks post immunization. This analysis revealed a greatly increased affinity of antigen-specific IgG for ALD vs. bolus vaccination, with a mean off-rate for antibodies elicited by ALD vaccination that was more than 10-fold lower than bolus at week 17 (**Fig. 6H**). Thus, ALD delivery substantially enhanced the evolution of antigen-specific GC B cells and subsequent serum antibody responses.

Finally, we sought to determine if the augmented GC responses promoted by ALD delivery would impact downstream development of long-lived plasma cells that home to the bone marrow and promote durable antibody production. As plasma cell development is particularly promoted following boosting, we compared bolus antigen with SMNP immunization as a prime only, a prime followed by a soluble boost 3 weeks later, or a single-shot prime-boost delivery obtained by co-injecting ALD30 and ALD100 particles loaded with equivalent doses of antigen. Animals were sacrificed for analysis of antigen-specific antibody secreting cells (ASCs) in the bone marrow at week 16. As expected, prime/boost bolus immunization elicited a much greater ASC population compared to prime-only bolus vaccination (**Fig. 6I**). However, single-shot ALD vaccination elicited a robust plasma cell response, ∼2-fold greater than the bolus prime/boost control (**Fig. 6I**). Thus, ALD vaccines may also have potential to improve the durability of humoral responses to vaccination.

## DISCUSSION

Vaccine delivery approaches that can prolong antigen delivery and program the timing of antigen release are of great interest for maximizing the priming and affinity maturation of rare protective B cell clones for HIV and other viral pathogens. In this study, we showed that ALD technology can enable both of these features of controlled vaccine kinetics. Notably, ALD particles carrying antigen alone without additional adjuvants elicited antibody and GC responses that were superior to bolus vaccination with a well studied saponin/TLR4 agonist adjuvant, SMNP, which we had previously shown to be more potent than a wide range of experimental and clinical adjuvants (*21, 53, 63*).

Based on the highly uniform coating thickness obtained in the atomic layer deposition process, it was expected that ALD particles might release vaccine antigen over a relatively narrow window of time once the alumina coatings dissolved *in vivo*. However, we found unexpectedly that antigen at ALD injection sites exhibited a period of very slow clearance determined by the number of alumina layers, followed by an extended period of more rapid antigen signal decay lasting ∼4 weeks. Several potential explanations for this slow-release behavior could be envisioned, including the formation of a fibrous capsule around the particle depot that could impact vaccine transport from the injection site or capture of a major pool of antigen by infiltrating immune cells. Interestingly, histological analysis of the injection site depots revealed that ALD particles induced no detectable fibrous capsule formation over at least 2 weeks, and that immune cell infiltration to the site was modest. Rather, the ALD particles appeared to show a pattern of release initiated from particles at the edge of the depot initially, followed by progression of particle uncoating from the outside of the depot to the interior. We hypothesize this pattern reflects the very low solubility of aluminum ions, making aluminum clearance at the edges of the depot a rate limiting step determining the time for ALD shell dissolution. Further work will be needed to test this idea.

The prolonged antigen release window achieved by ALD particle depots mimics “extended dosing” strategies where antigen is continuously delivered to draining lymph nodes over periods of a few weeks, which have been shown to impressively augment GC responses, increasing the number of unique B cell clones in GCs and promoting enhanced memory and plasma cell generation (*20, 60*). Mirroring findings from such extended dosing studies, we found that antigen was deposited on FDCs in the draining lymph nodes following ALD immunization, as well as robustly accumulated in dendritic cells. These patterns of antigen delivery correlated with enhanced expansion of affinity-matured B cells targeting HIV Env immunogens, which is a key goal for shepherding of humoral responses toward the development of broadly neutralizing antibodies. This provides a second important mechanism for tailoring the immune response in addition to the ability to program the timing of when antigen release initiates, a unique combination of features for this vaccine delivery technology.

In summary, ALD technology enables multiple critical elements of the immune response needed for the future development of vaccines against difficult-to-neutralize pathogens. ALD particles enable vaccine delivery kinetics to be tuned and drives potent humoral immune responses while avoiding overt inflammation, an ideal combination of properties for prophylactic vaccines.

## METHODS

### Protein production

N332-GT2 Env antigen and anti-CD45 VHH (clone G7) were expressed in FreeStyle 293F cells (Invitrogen, Waltham, MA). The VHH was purified by affinity chromatography using a GE HisTrap column while the antigen was purified in two steps, first by affinity chromatography using a GE HisTrap column followed by size-exclusion chromatography using a GE S200 column as previously described (*54*).

### Protein labelling

Antigen and VHH were fluorescently labeled with Alexa Fluor 647 and DyLight 405, respectively, using NHS ester chemistry. Briefly, proteins were incubated with fluorescent NHS esters (Lumiprobe, Cockeysville, MD and Thermo Fisher, Waltham, MA, respectively) at 6-fold molar excess of the protein in 0.2 M sodium bicarbonate buffer (pH 8.4) at 4 °C overnight at a final protein concentration of 2 mg/mL. Unbound dye was removed using Amicon Ultra-filters (Millipore Sigma, Burlington, MA). A 30 kDa cutoff filter was used for the antigen and a 3 kDa filter was utilized for the VHH. Proteins were filtered until the flow through was transparent. Concentration and degree of labeling were measured using a Nanodrop UV-vis spectroscopy instrument (Thermo Fisher, Waltham, MA). Labeled proteins were stored at 4 °C until use.

### SMNP production

SMNP adjuvant was produced as previously described with some modifications (*53*). Briefly, cholesterol (Avanti, Alabaster, AL), DPPC (Avanti, Alabaster, AL), MPLA (Avanti, Alabaster, AL), Quil-A saponin (Invivogen, San Diego, CA) were dissolved at a final concentration of 1 mg/mL, 0.5 mg/mL, 0.5 mg/mL, and 5 mg/mL, respectively, in 7.5% MEGA-10 detergent (Sigma Aldrich, St. Louis, MO) in Milli-Q water. The solution was passed through a 0.2 μm Supor syringe filter, diluted slowly 100-fold with PBS, and mixed gently overnight at 25 °C. Next, the solution was concentrated 50-fold by cross flow filtration (Thermo Fisher, Waltham, MA) and dialyzed for 4 days at 25 °C, against PBS using a 10 kDa MWCO dialysis cassette (Thermo Fisher, Waltham, MA) while changing dialysis buffer daily. The solution was sterilized using a 0.2 μm syringe filter and stored at 4 °C until use. SMNP concentration was determined using a cholesterol quantification kit (Sigma Aldrich, St. Louis, MO) as per the manufacturer’s instructions, and morphology was verified by negative-stain transmission electron microscopy.

### Lyophilization for screening of antigen stability in glassy polysaccharide formulations

Antigen was formulated at 50 μg/mL in formulations consisting of 9.5 w/v% endotoxin-free trehalose (Pfanstiehl, Waukegan, IL) and 10 mM or 50 mM L-histidine (Research Products International, Mt. Prospect, IL) with or without 2.5% hydroxyethyl starch (McKesson, Irving, TX), 40 mM ammonium acetate (JT Baker, Phillipsburg, NJ), and 0.2 w/w% Tween-20 (Thermo Fisher, Waltham, MA) at pH 5.5 or 6.5 according to **Table S1**. Hydroxyethyl starch was exchanged into 50 mM L-histidine by tangential flow filtration prior to use to remove salt. One mL aliquots of formulated antigen were filled into 3 mL Type II glass vials (Schott, Mainz, Germany) with 13-mm rubber butyl stoppers (DWK, Wertheim, Germany) inserted halfway. Formulated antigen was lyophilized using an FTS Systems LyoStar II lyophilizer (Warminster, PA). Filled vials were loaded onto shelves cooled to -1 °C and surrounded by a ring of unstoppered vials containing 1 mL water. The shelf temperature was decreased to -40 °C at an average rate of 2 °C/min to initiate freezing. Primary drying was initiated by decreasing the pressure to 60 mTorr. The temperature was increased to -20 °C at a rate of 1 °C/min then held at -20 °C for 24 hours. For secondary drying the temperature was raised to 40 °C at 1 °C/min and maintained at 40 °C. After 4 hours, the temperature was decreased to 4 °C and the chamber was backfilled with dry nitrogen to atmospheric pressure. The vials were fully stoppered and crimped and stored at 4 °C or 50 °C until further analysis. For assays, lyophilized samples were reconstituted to their original volume in sterile Milli-Q water.

### Spray-drying and alumina coating

Antigen was spray-dried as previously described (*64*). Briefly, antigen was formulated in 9.5 w/v% endotoxin-free trehalose (Pfanstiehl, Waukegan, IL), 2.5 % hydroxyethyl starch (B.O.C. Sciences, Shirley, NY), 50 mM L-histidine (Research Products International, Mt. Prospect, IL), 40 mM ammonium acetate (JT Baker, Phillipsburg, NJ) and 0.2 w/w% Tween-20 (Thermo Fisher, Waltham, MA) in Milli-Q water. All formulations were spray-dried in a Buchi B-290 Mini Spray Dryer (Buchi Labortechnik, Flawil, Switzerland) fitted with a two-fluid nozzle. Dehumidified air at an inlet temperature of 88°C and a flow rate of 32 m^3^/hr was used as a drying gas. Nitrogen gas was used in the atomizing nozzle at a flow rate of 414 L/hr, while the liquid sample was fed at 1 mL/min. These conditions yielded a spray-drier outlet temperature of less than 50 °C. Particles were collected in a high-performance cyclone separator. Water content was measured by Karl Fischer titration to be approximately 5%. Following spray-drying, powders were removed from the sample collector and aliquoted into 5 mL glass vials. Vials were placed on the shelf of an FTS Systems LyoStar II lyophilizer (Warminster, PA), where they were held overnight at 60 mTorr and 40 °C. Vials were then backfilled with dry nitrogen gas, sealed using butyl rubber stoppers and aluminum caps (DWK, Millville, NJ), and stored at 4 °C until use.

### ALD processing of spray-dried microspheres

Spray-dried powders were coated with 50–300 layers of alumina using alternating gas-phase injections of trimethylaluminum and water (**Fig. 1a**) in a custom-built fluidized bed atomic layer deposition reactor, as previously described (*64*). Zirconia/silicon spherical beads (0.1 mm OD) were mixed with the spray-dried microparticles at a 3:1 weight ratio prior to the ALD coating. The first 20 coating cycles were carried out with the beads present to reduce agglomeration of the initially cohesive and partially agglomerated polysaccharide microparticles until a conformal, less cohesive alumina layer was formed around the individual particles. The beads were then removed by sieving and coating cycles were continued until the desired layer thickness was achieved. The alumina content of the resulting powders was determined by weight after calcination to remove volatilizable excipients.

### Focused ion-beam scanning electron microscopy (FIB-SEM) analysis of ALD layer

ALD50, ALD100, and ALD200 microparticles were analyzed at the University of Colorado Boulder Colorado Shared Instrumentation in Nanofabrication and Characterization Facility (COSINC). Powders were adhered to a pin mount with dual-sided tape. Samples were coated with a Leica EM ACE600 High Vacuum Sputter Coater (Leica Microsystems, Boston, MA) with an 8.97 nm Pt layer, at a rotating 15° tilt. A FEI Nova 600i Dual Beam with a Ga Focused Ion Beam and Field Emission SEM (Thermo Fisher, Waltham, MA) was used to create a cross-section of particles with additional localized Pt deposition using a 28 pA FIB with a z coordinate set to 0.2 nm. Images were taken at a 52° tilt, and the coating layer thickness was analyzed using ImageJ 1.54g (*65*) .

### Antigenicity profiling

Antigenicity profiling was performed on antigen recovered from either spray-dried or ALD particles. Spray-dried particles were directly reconstituted in PBS. To recover antigen from coated ALD particles, samples were mechanically disrupted by vortexing with a stir bar for eight 1-minute intervals. Disrupted ALD particles were then resuspended in an alumina dissolution buffer containing 90.5 mM sodium phosphate, 20.5 mM citric acid, and 0.06% Polysorbate 80 at pH 6.6 for 1 hour at 4 °C under agitation. Subsequently, samples were centrifuged to pellet residual alumina and the supernatant containing the recovered antigen was collected. Recovered antigen was stored at 4 °C until further use.

For ELISA, high-binding half-area polystyrene plates (Corning, NY) were coated overnight at 4 °C with 2 µg/mL *Galanthus nivalis* lectin (Invitrogen, Waltham, MA) in PBS. The following day, plates were washed three times with 0.05% Tween-20 in PBS (Millipore Sigma, Burlington, MA) and blocked with 2% bovine serum albumin (Invitrogen, Waltham, MA) in PBS (blocking buffer) at 25 °C to prevent nonspecific binding. All subsequent incubations were performed at 25 °C. Plates were then incubated with 2 µg/mL of recovered antigen samples in blocking buffer for 2 hr. Plates were washed three times, and anti-Env antibodies were added, with the top-row concentration at 0.3 µg/mL, followed by 4-fold serial dilutions. After two hours, plates were washed three times, and incubated with a goat anti-human HRP secondary antibody (Bio-Rad, Hercules, CA) in blocking buffer for 1 hour. Plates were then washed 5 times and developed for 5 min with 50 µL of tetramethylbenzidine substrate. The reaction was stopped with 25 µL of 2 N sulfuric acid (Ricca Chemical, Arlington, TX), and absorbance was measured at 450/570 nm using a plate reader.

### Bone marrow dendritic cell culture

Bone marrow–derived dendritic cells (BMDCs) were generated from femurs and tibias of C57BL/6 mice. Briefly, bone marrow was flushed with sterile RPMI-1640 (Gibco, Waltham, MA) using a 25-gauge needle, passed through a 70 µm cell strainer, and pelleted by centrifugation. Red blood cells were lysed using ACK lysis buffer, and cells were washed and resuspended in complete RPMI-1640 (Gibco, Waltham, MA) supplemented with 10% fetal bovine serum (Gibco, Waltham, MA), 1% penicillin–streptomycin (Gibco, Waltham, MA), 2 mM L-glutamine (Gibco, Waltham, MA), 1 mM sodium pyruvate (Gibco, Waltham, MA), and 50 µM β-mercaptoethanol (Gibco, Waltham, MA). Cells were counted and plated in non-adherent petri dishes at a density of 2 million cells in 10 mL of medium and cultured at 37 °C with 5% CO₂ in the presence of 20 ng/mL GM-CSF and 20 ng/mL IL-4 (PeproTech, Cranbury, NJ). On day 3, half of the culture medium was gently removed and replaced with fresh complete medium containing 20 ng/mL GM-CSF and 20 ng/mL IL-4. On day 7, the medium was refreshed again in the same manner. Loosely adherent and non-adherent immature BMDCs were harvested on days 8 or 9 for downstream experiments.

### ALD particle uptake by RAW 264.7 cells

RAW 264.7 cells (ATTC TIB-71) were maintained in Dulbecco’s Modified Eagle Medium (DMEM), 10 % (v/v) fetal bovine serum (FBS), and 1% (v/v) antibiotic / antimycotic solution. Cells were maintained in 5% CO_2_ and at 37°C. Cells were routinely subcultured every 3-4 days.

For microscopy analysis, fluorescent particles containing Alexa Fluor 647-labeled antigen and 30 alumina layers were formulated at 0.8 µg/mg powder. RAW 264.7 cells were seeded at 4000 cells/well in a culture-treated 96-well plate (PerkinElmer, Shelton, CT) in DMEM with 10% FBS and incubated overnight. The cells were then incubated with 100 ng/mL LPS for 2 hours prior to the introduction of 200 µL of a 10 µg/mL fluorescent ALD30 suspension in 10% FBS DMEM. After a 24-hour incubation with the particles, the cell membrane was stained with DiL, and nuclei were stained with Hoechst 33342. Lastly, cells were fixed with 4% paraformaldehyde and stored protected from light until imaging on a Revvity PerkinElmer Opera Phenix using the 40x water immersion objective. The imaging work was performed at the BioFrontiers Institute’s Advanced Light Microscopy Core (RRID: SCR_018302, NIH grant 1S10OD025072).

For flow cytometry analysis, fluorescent particles containing a calcein fluorescent probe and 50 alumina layers were formulated at 1.2 µg calcein/mg powder. RAW 264.7 cells were seeded at 10^5^ cells/well in 12-well culture-treated plates. Unactivated control groups were seeded in the maintenance media described above while activated groups were seeded in DMEM supplemented with 5% FBS, 1% antibiotic/antimycotic solution and 100 ng/mL LPS for 22 hours prior to introduction of the calcein particles. Particles were suspended at the desired concentration in the activation media formulation. At 22 hours the media was replaced with particle-containing media and allowed to incubate for 2 hours. After 2 hours, wells were washed with PBS 2-3 times to remove excess particles and cells were detached from plates by incubating with 0.25% trypsin-EDTA for 5 minutes, followed by gentle pipetting. Activation towards M1-like phenotype was confirmed using CD86 monoclonal antibody (Thermo Fisher, Waltham, MA) (captured using 450/40 filter set with 405 nm excitation). Particle uptake was analyzed based on fluorescence read out in a 530/30 filter set following excitation with a 488 nm laser. Measurements were carried out on either a BD FACSCelesta or BD Accuri flow cytometer. Analysis was completed using FlowJo at Boulder Flow Cytometry Shared Core Facility (RRID: SCR_019309).

### ALD uptake and activation in BMDC

Bone marrow–derived dendritic cells (BMDCs) were seeded in 24-well tissue culture plates at a density of 0.5 million cells/mL and maintained at 37 °C in a humidified incubator with 5% CO₂. ALD particles encapsulating Alexa Fluor 647-labeled antigen were resuspended directly in complete culture media and added to cells at a final concentration of 0.5 mg particles/mL (total particle mass). Lipopolysaccharide (LPS from E. coli O11:B4; Millipore Sigma) was used at 10 ng/mL as a positive control for BMDC activation. Cells were incubated with treatments for 24 hrs. To assess cell viability and activation, cells were harvested and incubated with 1X Zombie Aqua fixable viability dye (BioLegend, San Diego, CA) in PBS. Fc receptors were blocked using mouse TruStain FcX^TM^ PLUS (BioLegend, San Diego, CA) in FACS buffer consisting of PBS supplemented with 2% FBS and 5 mM EDTA. Cells were subsequently stained with fluorophore-conjugated antibodies (BioLegend, San Diego, CA) against surface markers. BMDCs were identified as CD45⁺ CD11b⁺ CD11c⁺ F4/80⁻ cells. Particle uptake and activation were assessed by flow cytometry, with CD80 and CD86 used as activation markers. Data were acquired on a Cytek Aurora spectral flow cytometer and analyzed using FlowJo v10 (BD Biosciences, Franklin Lakes, NJ). Caspase-1 activity and ATP release were measured in a separate set of cells using the Caspase-Glo^®^ 1 Inflammasome Assay (Promega, Madison, WI) and the RealTime-Glo™ Extracellular ATP Assay (Promega, Madison, WI), respectively, as per the manufacturer’s instructions.

### ALD pharmacokinetics *in vivo*

ALD particles (∼1.9×10^8^) of varying coating thicknesses encapsulating 10 μg Alexa Fluor 647-labeled antigen were subcutaneously administered in female C57BL/6 mice near the tail base. A liquid bolus dose of 10 μg Alexa Fluor 647-labeled antigen and 5 μg SMNP adjuvant were used as a reference. Mice were longitudinally imaged using an Ami HTX imaging system (Bruker, Billerica, MA) at ex/em: 640/690 nm. Total radiant efficiency (p/s)/(μW/cm^2^) was calculated using region of interest analysis using Spectral Aura software and normalized to fluorescence 6 hrs post injection.

### Antibody ELISA

High-binding half-area polystyrene plates (Corning, NY) were coated overnight at 4 °C with 2 µg/mL streptavidin (Invitrogen, Waltham, MA). The following day, plates were washed three times with 0.05% Tween-20 in PBS (Millipore Sigma, Burlington, MA) and blocked with 2% bovine serum albumin (Invitrogen, Waltham, MA) in PBS (blocking buffer) at 25 °C to prevent nonspecific binding. All subsequent incubations were performed at 25 °C. Plates were then coated with 2 µg/mL biotinylated antigen in blocking buffer for 2 hours. Serum samples were serially diluted 4-fold, starting at a 1:50 dilution, and added to the washed plates. After 2 hours, the plates were washed three times and incubated with a goat anti-mouse HRP secondary detection antibody (Bio-Rad, Hercules, CA) in blocking buffer for an hour. Plates were then washed five times, 50 µL of tetramethylbenzidine substrate was added, and the color was allowed to develop for 5 min. The reaction was stopped with 25 µL of 2 N sulfuric acid (Ricca Chemical, Arlington, TX), and absorbance was measured at 450/570 nm using a plate reader.

### Fluorescent antigen uptake *in vivo*

Particles containing 10 μg Alexa Fluor 647-labeled antigen encapsulated in ALD50 were administered subcutaneously near the tail base of female C57BL/6 mice. As a comparator, mice received 10 µg of labeled antigen adjuvanted with 5 µg of SMNP as a liquid bolus. Mice were euthanized at indicated time points post-injection. Skin from the injection site and draining lymph nodes (inguinal, axillary, and brachial) were harvested for downstream analysis.

Harvested skin was digested in RPMI-1640 containing 0.5 mg/mL collagenase D (Millipore Sigma, Burlington, MA) , 0.5 mg/mL DNase I (Millipore Sigma, Burlington, MA), and 5 mM CaCl₂ for 2 hours at 37 °C under agitation. Following digestion, tissue was passed through a 70 µm cell strainer, single cell suspensions were pelleted by centrifugation, and red blood cells were lysed using ACK lysis buffer (Gibco, Waltham, MA). Cells were then resuspended in FACS buffer consisting of 2% FBS in PBS with 5 mM EDTA. Lymph nodes (axillary, brachial, and inguinal) were pooled and mechanically disrupted before being digested in RPMI-1640 containing 0.1 mg/mL collagenase D, 0.1 mg/mL DNase I, and 5 mM CaCl₂ for 20 min at 37 °C under agitation. Digested lymph node suspensions were passed through a 70 µm cell strainer, pelleted by centrifugation, and resuspended in FACS buffer.

Single cell suspensions from both skin and lymph nodes were pelleted and stained with 1X Zombie Aqua viability dye (Biolegend, San Diego, CA) in PBS for 15 min at 25°C. Cells were then incubated with mouse TruStain FcX^TM^ PLUS (Biolegend, San Diego, CA) for 20 min at 4 °C to block Fc receptors. Following Fc block, cells were stained with a fluorescent antibody cocktail for 30 min at 4 °C. After staining, cells were resuspended in FACS buffer. Lymph nodes were topped with Precision count beads (10 μm, Biolegend, San Diego, CA), and skin was topped with Accucount beads (15 μm, Spherotech, Lake Forest, IL). Cells were analyzed on a Cytek Aurora spectral flow analyzer (Cytek Biosciences, Fremont, CA). In some experiments, cells were analyzed on a CytPix Attune flow cytometer (Thermo Fisher, Waltham, MA) to acquire brightfield images and determine whether cells were particle-associated or had internalized soluble antigen. Data were processed in Flowjo v10 (Waters Corp., Milford, MA).

### Immunofluorescence microscopy

Tissues were fixed in 0.075 M L-lysine, 0.0375 M sodium phosphate, 0.01 M sodium periodate, and 1% formaldehyde (all Millipore Sigma, Burlington, MA) for 4 hours at 25 °C. Lymph nodes were cryoprotected by sequential immersion in 15% and 30% sucrose in PBS, embedded in OCT (Sakura Finetek, Torrance, CA), frozen in liquid nitrogen-chilled isopentane, and cryosectioned at 12 µm on a Leica CM1950 (Leica Biosystems, Deer Park, IL) onto Superfrost Plus slides (Fisher Scientific, Waltham, MA). Sections were stored at −20 °C. Injection sites were processed identically but sectioned at 20 µm, or alternatively sectioned at 150 µm on a vibrating microtome (Precisionary Instruments, Ashland, MA) and stored in PBS.

For staining, sections were rehydrated in 0.05% Tween-20 in PBS (wash buffer, 5 min), blocked (5% FBS in PBS, 1 h, 25 °C), and permeabilized (2% Triton X-100 in PBS, 30 min). Lymph node sections were incubated overnight at 4 °C with BV421 anti-CD35 (clone 8C12, BD Biosciences, Franklin Lakes, NJ), Alexa Fluor 594 anti-B220 (clone RA3-6B2, Biolegend, San Diego, CA), and Alexa Fluor 647 anti-Env (clone BG18) diluted in staining buffer (0.2% Triton X-100 and 5% FBS in PBS). Injection site sections were incubated with DyLight 405 anti-CD45 VHH (clone G7) overnight at 4 °C in staining buffer. All fluorescent reagents were added at a final concentration of 2 μg/mL. After three washes, cryosections were mounted with ProLong Diamond antifade medium (Thermo Fisher, Waltham, MA) and microtome sections with Slow Fade Glass medium (Thermo Fisher, Waltham, MA) with a 0.1 mm spacer (SUNJin Labs, Hsinchu City, Taiwan), both under #1.5 coverslips (Electron Microscopy Sciences, Hatfield, PA). Images were acquired on a Zeiss LSM 780 (Carl Zeiss Microscopy, White Plains, NY) and analyzed in Fiji (*66*).

Immune cell infiltration was quantified by measuring CD45 fluorescence intensity within manually defined regions of interest (ROIs) inside the depot and in the surrounding tissue. Relative immune cell intensity was calculated as:

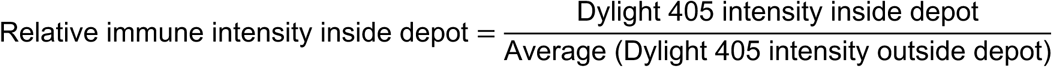

Antigen–particle co-retention was assessed by quantifying colocalization between the calcein and antigen channels using the coloc2 plugin in Fiji. ROIs were drawn within the depot region, and colocalization was reported as Manders’ overlap coefficient, restricted to images passing a Costes randomization threshold (p > 0.95).

### Germinal center responses

Female C57BL/6 mice were subcutaneously immunized with 10 µg antigen delivered either as a liquid bolus dose with 5 μg SMNP adjuvant in PBS or encapsulated within ALD50 particles suspended in triglyceride oil. Mice were euthanized at the specified endpoints, and lymph nodes (inguinal, axillary, and brachial) were harvested for analysis. Lymph nodes were mechanically disrupted and passed through a 70 µm cell strainer to generate single-cell suspensions. Next, cells were pelleted and resuspended in 1X Zombie Aqua viability dye (Biolegend, San Diego, CA) for 15 min at 25°C. Following viability staining, cells were centrifuged and resuspended in mouse TruStain FcX^TM^ PLUS (1:100, Biolegend, San Diego, CA) for 20 min at 4°C to block Fc receptors. Tetramers were formulated by incubating fluorescent streptavidin (BV421 and PE) with biotinylated antigen at a 3:1 streptavidin: antigen mass ratio for at least one hour at 25°C. After Fc receptor blockade, cells were pelleted and stained with 1 μg antigen tetramer each in 50 μL FACS buffer for 45 min. After tetramer staining, cells were spun down and resuspended in an antibody cocktail for 30 min at 4 °C. Next, cells were spun down, resuspended in FACS buffer, and supplemented with Precision count beads (10 μm, Biolegend, San Diego, CA). Cells were analyzed using a Cytek Aurora spectral flow cytometer (Cytek Biosciences, Fremont, CA). Data were processed in Flowjo v10 (Waters Corp., Milford, MA).

### Bio-layer interferometry (BLI)

Bio-layer interferometry (BLI) was performed on an Octet R8 instrument (Sartorius, Bohemia, NY) at 25 °C using streptavidin (SA) biosensors. Sensors were hydrated in PBS supplemented with 0.05% (v/v) Tween-20 prior to use. Biotinylated antigen was immobilized onto SA sensors at 25 µg/mL for 30 sec to achieve a target loading response of ∼1.2 nm, followed by a 60 sec baseline in buffer. Serum samples were diluted 1:50 in PBS + 0.05% Tween-20 and applied to antigen-loaded sensors for an association phase of 600 sec, followed by dissociation in buffer for 300 sec. Dissociation rate constants (k_off_) were determined by fitting the dissociation phase using the Octet data analysis software.

### Legendplex

Serum cytokine and chemokine levels (IFN-γ, CXCL1, TNF-α, CCL2, IL-12p70, CCL5, IL-1β, CXCL10, GM-CSF, IL-10, IFN-β, IFN-α, and IL-6) were quantified using the LEGENDplex™ Mouse Anti-Virus Response Panel (BioLegend, San Diego, CA) according to the manufacturer’s protocol. Samples were acquired on a Cytek Aurora and data were analyzed using FlowJo v10.

### Bone marrow ELISPOT

Antigen-specific plasma cells in bone marrow were quantified using the ELISpot Flex: Mouse IgG (HRP) kit (Mabtech, Cincinnati, OH ) according to the manufacturer’s instructions with minor modifications. Bone marrow cells were isolated from the femurs of C57BL/6 mice as previously described and resuspended in RPMI-1640 supplemented with 10% fetal bovine serum (cell media). PVDF 96-well ELISPOT plates (51-2447KC, BD Biosciences, Franklin Lakes, NJ) were pre-wetted with 70% ethanol for 90 sec, washed with PBS, and coated overnight at 4 °C with 10 µg/mL goat anti-mouse IgG capture antibody. Plates were washed five times with PBS and blocked with cell media for two hours at 25°C. Next, 100 μL of bone marrow cells were added to the plates at a concentration of 1 million cells/mL and incubated at 37 °C in a humidified incubator with 5% CO₂ for 16 hours to allow antibody secretion and capture. After incubation, cells were removed, and plates were washed five times with PBS between each subsequent step. Biotinylated antigen was used as the detection reagent at a concentration of 2 µg/mL. Plates were washed five times with PBS and incubated with streptavidin-HRP at a 1:100 dilution. Following an additional five PBS washes, spots were developed using AEC substrate (BD Biosciences, Franklin Lakes, NJ) for 5 min at 25 °C. The reaction was stopped by rinsing thoroughly with water, and plates were allowed to dry overnight. Spots were quantified using a ImmunoSpot ELISPOT reader (CTL, Shaker Heights, OH).

## ACKNOWLEDGMENTS

This work was supported by the Gates Foundation (IN-050230 to DJI, INV-064813 to DJI, TWR and RLG) and the NIH (UM1AI144462 to DJI, AI192197 to DJI and TWR). DJI is an investigator of the Howard Hughes Medical Institute. Tomoko Borsa at the University of Colorado, Boulder COSINC facility obtained FIB-SEM images of alumina layer thicknesses.

## AUTHOR CONTRIBUTIONS

Conceptualization: DJI, TWR, RLG, NC

Methodology: NC, DJI, TWR, RLG

Investigation: NC, HJC, EVP, JY, AWL, AAL, AAW, HS, AW, MMM, JBG, MBM, AMR, ESL, UP, MOP, CAW, HHF

Visualization: NC, DJI

Statistical Analysis: NC

Funding acquisition: DJI, TWR, RLG

Supervision: DJI, TWR, RLG

Writing – original draft: DJI, NC, TWR, RLG

Writing – review & editing: All authors

## COMPETING INTERESTS

TWR, HJC, HHF, and RLG are inventors on patents and patent applications related to ALD vaccines, which are licensed to Vitrivax Inc. HHF is employed by VitriVax, Inc. DJI is an inventor on patents related to SMNP adjuvant.

## Notes

### Summary of Updates

Supplemental Figure 3D added to indicate that ALD doesn't trigger inflammatory cytokines.

## REFERENCES

1. A. J. Shattock, H. C. Johnson, S. Y. Sim, A. Carter, P. Lambach, R. C. W. Hutubessy, K. M. Thompson, K. Badizadegan, B. Lambert, M. J. Ferrari, M. Jit, H. Fu, S. P. Silal, R. A. Hounsell, R. G. White, J. F. Mosser, K. A. M. Gaythorpe, C. L. Trotter, A. Lindstrand, K. L. O’Brien, N. Bar-Zeev, Contribution of vaccination to improved survival and health: modelling 50 years of the Expanded Programme on Immunization. Lancet 403, 2307–2316 (2024).

2. C. M. C. Rodrigues, S. A. Plotkin, Impact of Vaccines; Health, Economic and Social Perspectives. Front Microbiol 11, 1526 (2020).

3. G. D. Victora, M. C. Nussenzweig, Germinal Centers. Annu. Rev. Immunol. 40, 1–30 (2022).

4. I. J. Amanna, N. E. Carlson, M. K. Slifka, Duration of Humoral Immunity to Common Viral and Vaccine Antigens. N. Engl. J. Med. 357, 1903–1915 (2007).

5. C. W. Tan, S. A. Valkenburg, L. L. M. Poon, L.-F. Wang, Broad-spectrum pan-genus and pan-family virus vaccines. Cell Host Microbe 31, 902–916 (2023).

6. J.-Y. Park, H.-M. Lee, Optimizing Humoral Immunity for Durable and Broad Protection in Flavivirus Vaccines. Vaccines 13, 1182 (2025).

7. C.-J. Wei, M. C. Crank, J. Shiver, B. S. Graham, J. R. Mascola, G. J. Nabel, Next-generation influenza vaccines: opportunities and challenges. Nat Rev Drug Discov 19, 239–252 (2020).

8. L. E. Rosen, M. A. Tortorici, A. D. Marco, D. Pinto, W. B. Foreman, A. L. Taylor, Y.-J. Park, D. Bohan, T. Rietz, J. M. Errico, K. Hauser, H. V. Dang, J. W. Chartron, M. Giurdanella, G. Cusumano, C. Saliba, F. Zatta, K. R. Sprouse, A. Addetia, S. K. Zepeda, J. Brown, J. Lee, E. Dellota, A. Rajesh, J. Noack, Q. Tao, Y. DaCosta, B. Tsu, R. Acosta, S. Subramanian, G. D. de Melo, L. Kergoat, I. Zhang, Z. Liu, B. Guarino, M. A. Schmid, G. Schnell, J. L. Miller, F. A. Lempp, N. Czudnochowski, E. Cameroni, S. P. J. Whelan, H. Bourhy, L. A. Purcell, F. Benigni, J. di Iulio, M. S. Pizzuto, A. Lanzavecchia, A. Telenti, G. Snell, D. Corti, D. Veesler, T. N. Starr, A potent pan-sarbecovirus neutralizing antibody resilient to epitope diversification. Cell 187, 7196–7213.e26 (2024).

9. D. Stadlbauer, X. Zhu, M. McMahon, J. S. Turner, T. J. Wohlbold, A. J. Schmitz, S. Strohmeier, W. Yu, R. Nachbagauer, P. A. Mudd, I. A. Wilson, A. H. Ellebedy, F. Krammer, Broadly protective human antibodies that target the active site of influenza virus neuraminidase. Science 366, 499–504 (2019).

10. C. Havenar-Daughton, A. Sarkar, D. W. Kulp, L. Toy, X. Hu, I. Deresa, O. Kalyuzhniy, K. Kaushik, A. A. Upadhyay, S. Menis, E. Landais, L. Cao, J. K. Diedrich, S. Kumar, T. Schiffner, S. M. Reiss, G. Seumois, J. R. Yates, J. C. Paulson, S. E. Bosinger, I. A. Wilson, W. R. Schief, S. Crotty, The human naive B cell repertoire contains distinct subclasses for a germline-targeting HIV-1 vaccine immunogen. Science Translational Medicine 10, eaat0381 (2018).

11. J. M. Steichen, Y.-C. Lin, C. Havenar-Daughton, S. Pecetta, G. Ozorowski, J. R. Willis, L. Toy, D. Sok, A. Liguori, S. Kratochvil, J. L. Torres, O. Kalyuzhniy, E. Melzi, D. W. Kulp, S. Raemisch, X. Hu, S. M. Bernard, E. Georgeson, N. Phelps, Y. Adachi, M. Kubitz, E. Landais, J. Umotoy, A. Robinson, B. Briney, I. A. Wilson, D. R. Burton, A. B. Ward, S. Crotty, F. D. Batista, W. R. Schief, A generalized HIV vaccine design strategy for priming of broadly neutralizing antibody responses. Science 366, eaax4380 (2019).

12. R. W. Sanders, J. P. Moore, Progress on priming HIV-1 immunity. Science 384, 738–739 (2024).

13. B. F. Haynes, K. Wiehe, P. Borrow, K. O. Saunders, B. Korber, K. Wagh, A. J. McMichael, G. Kelsoe, B. H. Hahn, F. Alt, G. M. Shaw, Strategies for HIV-1 vaccines that induce broadly neutralizing antibodies. Nat. Rev. Immunol. 23, 142–158 (2023).

14. X. Chen, T. Zhou, S. D. Schmidt, H. Duan, C. Cheng, G.-Y. Chuang, Y. Gu, M. K. Louder, B. C. Lin, C.-H. Shen, Z. Sheng, M. X. Zheng, N. A. Doria-Rose, M. G. Joyce, L. Shapiro, M. Tian, F. W. Alt, P. D. Kwong, J. R. Mascola, Vaccination induces maturation in a mouse model of diverse unmutated VRC01-class precursors to HIV-neutralizing antibodies with >50% breadth. Immunity 54, 324–339.e8 (2021).

15. A. Escolano, J. M. Steichen, P. Dosenovic, D. W. Kulp, J. Golijanin, D. Sok, N. T. Freund, A. D. Gitlin, T. Oliveira, T. Araki, S. Lowe, S. T. Chen, J. Heinemann, K.-H. Yao, E. Georgeson, K. L. Saye-Francisco, A. Gazumyan, Y. Adachi, M. Kubitz, D. R. Burton, W. R. Schief, M. C. Nussenzweig, Sequential Immunization Elicits Broadly Neutralizing Anti-HIV-1 Antibodies in Ig Knockin Mice. Cell 166, 1445–1458.e12 (2016).

16. C. A. Cottrell, X. Hu, J. H. Lee, P. Skog, S. Luo, C. T. Flynn, K. R. McKenney, J. Hurtado, O. Kalyuzhniy, A. Liguori, J. R. Willis, E. Landais, S. Raemisch, X. Chen, S. Baboo, S. Himansu, J. K. Diedrich, H. Duan, C. Cheng, T. Schiffner, D. L. V. Bader, D. W. Kulp, R. Tingle, E. Georgeson, S. Eskandarzadeh, N. Alavi, D. Lu, T. Sincomb, M. Kubitz, T.-M. Mullen, J. R. YatesIII, J. C. Paulson, J. R. Mascola, F. W. Alt, B. Briney, D. Sok, W. R. Schief, Heterologous prime-boost vaccination drives early maturation of HIV broadly neutralizing antibody precursors in humanized mice. Sci. Transl. Med. 16, eadn0223 (2024).

17. K. Wiehe, K. O. Saunders, V. Stalls, D. W. Cain, S. Venkatayogi, J. S. M. Beem, M. Berry, T. Evangelous, R. Henderson, B. Hora, S.-M. Xia, C. Jiang, A. Newman, C. Bowman, X. Lu, M. E. Bryan, J. Bal, A. Sanzone, H. Chen, A. Eaton, M. A. Tomai, C. B. Fox, Y. K. Tam, C. Barbosa, M. Bonsignori, H. Muramatsu, S. M. Alam, D. C. Montefiori, W. B. Williams, N. Pardi, M. Tian, D. Weissman, F. W. Alt, P. Acharya, B. F. Haynes, Mutation-guided vaccine design: A process for developing boosting immunogens for HIV broadly neutralizing antibody induction. Cell Host Microbe 32, 693–709.e7 (2024).

18. J. M. Steichen, P. J. Madden, C. T. Flynn, S. Phulera, M. Shil, O. Kalyuzhniy, A. Liguori, C. Kifude, L. M. Sewall, C. A. Cottrell, K. M. Ma, S. Baboo, J. K. Diedrich, K. McKenney, A. C. deCamp, D. G. Carnathan, I. Phung, P. Ramezani-Rad, E. Marina-Zárate, B. Freeman, Z. Xie, J. H. Lee, T. Sincomb, N. Phelps, D. Lu, D. Goodwin, R. Tingle, Y. Adachi, N. Alavi, J. Tran, A. S. Tran, A. Nascimento, C. Sovie, D. L. V. Bader, H. Voic, X. Zhou, G. Pixton, A. Walsh, M. B. Melo, T. Schiffner, F. D. Batista, D. R. Burton, D. J. Irvine, J. C. Paulson, J. R. Yates, G. Ozorowski, A. B. Ward, G. Silvestri, S. Crotty, W. R. Schief, Vaccination elicits HIV broadly neutralizing antibodies in primates. Nature , 1–3 (2026).

19. J. H. Lee, H. J. Sutton, C. A. Cottrell, I. Phung, G. Ozorowski, L. M. Sewall, R. Nedellec, C. Nakao, M. Silva, S. T. Richey, J. L. Torres, W.-H. Lee, E. Georgeson, M. Kubitz, S. Hodges, T.-M. Mullen, Y. Adachi, K. M. Cirelli, A. Kaur, C. Allers, M. Fahlberg, B. F. Grasperge, J. P. Dufour, F. Schiro, P. P. Aye, O. Kalyuzhniy, A. Liguori, D. G. Carnathan, G. Silvestri, X. Shen, D. C. Montefiori, R. S. Veazey, A. B. Ward, L. Hangartner, D. R. Burton, D. J. Irvine, W. R. Schief, S. Crotty, Long-primed germinal centres with enduring affinity maturation and clonal migration. Nature 609, 998–1004 (2022).

20. K. M. Cirelli, D. G. Carnathan, B. Nogal, J. T. Martin, O. L. Rodriguez, A. A. Upadhyay, C. A. Enemuo, E. H. Gebru, Y. Choe, F. Viviano, C. Nakao, M. G. Pauthner, S. Reiss, C. A. Cottrell, M. L. Smith, R. Bastidas, W. Gibson, A. N. Wolabaugh, M. B. Melo, B. Cossette, V. Kumar, N. B. Patel, T. Tokatlian, S. Menis, D. W. Kulp, D. R. Burton, B. Murrell, W. R. Schief, S. E. Bosinger, A. B. Ward, C. T. Watson, G. Silvestri, D. J. Irvine, S. Crotty, Slow Delivery Immunization Enhances HIV Neutralizing Antibody and Germinal Center Responses via Modulation of Immunodominance. Cell 177, 1153–1171.e28 (2019).

21. P. J. Madden, E. Marina-Zárate, K. A. Rodrigues, J. M. Steichen, M. Shil, K. Ni, K. K. Michaels, L. Maiorino, A. A. Upadhyay, S. Saha, A. Pradhan, O. Kalyuzhiny, A. Liguori, P. G. Lopez, I. Phung, C. Flynn, A. Zhou, M. B. Melo, A. Lemnios, N. Phelps, E. Georgeson, N. Alavi, M. Kubitz, D. Lu, S. Eskandarzadeh, A. Metz, O. L. Rodriguez, K. Shields, S. Schultze, M. L. Smith, B. S. Healy, D. Lim, V. R. Lewis, E. Ben-Akiva, W. Pinney, J. Gregory, S. Xiao, D. G. Carnathan, S. P. Kasturi, C. T. Watson, S. E. Bosinger, G. Silvestri, W. R. Schief, D. J. Irvine, S. Crotty, Diverse priming outcomes under conditions of very rare precursor B cells. Immunity 58, 997–1014.e11 (2025).

22. J. M. Steichen, I. Phung, E. Salcedo, G. Ozorowski, J. R. Willis, S. Baboo, A. Liguori, C. A. Cottrell, J. L. Torres, P. J. Madden, K. M. Ma, H. J. Sutton, J. H. Lee, O. Kalyuzhniy, J. D. Allen, O. L. Rodriguez, Y. Adachi, T.-M. Mullen, E. Georgeson, M. Kubitz, A. Burns, S. Barman, R. Mopuri, A. Metz, T. K. Altheide, J. K. Diedrich, S. Saha, K. Shields, S. E. Schultze, M. L. Smith, T. Schiffner, D. R. Burton, C. T. Watson, S. E. Bosinger, M. Crispin, J. R. YatesIII, J. C. Paulson, A. B. Ward, D. Sok, S. Crotty, W. R. Schief, Vaccine priming of rare HIV broadly neutralizing antibody precursors in nonhuman primates. Science 384, eadj8321 (2024).

23. J. H. Eldridge, J. K. Staas, J. A. Meulbroek, T. R. Tice, R. M. Gilley, Biodegradable and biocompatible poly(DL-lactide-co-glycolide) microspheres as an adjuvant for staphylococcal enterotoxin B toxoid which enhances the level of toxin-neutralizing antibodies. Infect. Immun. 59, 2978–2986 (1991).

24. J. H. Eldridge, R. M. Gilley, J. K. Staas, Z. Moldoveanu, J. A. Meulbroek, T. R. Tice, New Strategies for Oral Immunization, International Symposium at the University of Alabama at Birmingham and Molecular Engineering Associates, Inc. Birmingham, AL, USA, March 21–22, 1988. Curr. Top. Microbiol. Immunol. 146, 59–66 (1989).

25. D. T. O’Hagan, D. Rahman, J. P. McGee, H. Jeffery, M. C. Davies, P. Williams, S. S. Davis, S. J. Challacombe, Biodegradable microparticles as controlled release antigen delivery systems. Immunology 73, 239–242 (1991).

26. J. Kohn, S. M. Niemi, E. C. Albert, J. C. Murphy, R. Langer, J. G. Fox, Single-step immunization using a controlled release, biodegradable polymer with sustained adjuvant activity. J. Immunol. Methods 95, 31–38 (1986).

27. J. L. Cleland, A. Lim, L. Barrón, E. T. Duenas, M. F. Powell, Development of a single-shot subunit vaccine for HIV-1: Part 4. Optimizing microencapsulation and pulsatile release of MN rgp120 from biodegradable microspheres. J. Control. Release 47, 135–150 (2010).

28. F. A. Sharp, D. Ruane, B. Claass, E. Creagh, J. Harris, P. Malyala, M. Singh, D. T. O’Hagan, V. Pétrilli, J. Tschopp, L. A. J. O’Neill, E. C. Lavelle, Uptake of particulate vaccine adjuvants by dendritic cells activates the NALP3 inflammasome. Proceedings of the National Academy of Sciences 106, 870–875 (2009).

29. G. F. Kersten, D. Donders, A. Akkermans, E. C. Beuvery, Single shot with tetanus toxoid in biodegradable microspheres protects mice despite acid-induced denaturation of the antigen. Vaccine 14, 1627–1632 (1996).

30. P. Malyala, D. T. O’Hagan, Immunopotentiators in Modern Vaccines (Second Edition). , 231–248 (2017).

31. B. A. Bailey, L. J. Ochyl, S. P. Schwendeman, J. J. Moon, Toward a Single-Dose Vaccination Strategy with Self-Encapsulating PLGA Microspheres. Adv Healthc Mater 6, 1601418 (2017).

32. X. Xi, T. Ye, S. Wang, X. Na, J. Wang, S. Qing, X. Gao, C. Wang, F. Li, W. Wei, G. Ma, Self-healing microcapsules synergetically modulate immunization microenvironments for potent cancer vaccination. Sci Adv 6, eaay7735 (2020).

33. S. Y. Tzeng, K. J. McHugh, A. M. Behrens, S. Rose, J. L. Sugarman, S. Ferber, R. Langer, A. Jaklenec, Stabilized single-injection inactivated polio vaccine elicits a strong neutralizing immune response. Proc National Acad Sci 115, E5269–E5278 (2018).

34. S. Y. Tzeng, R. Guarecuco, K. J. McHugh, S. Rose, E. M. Rosenberg, Y. Zeng, R. Langer, A. Jaklenec, Thermostabilization of inactivated polio vaccine in PLGA-based microspheres for pulsatile release. Journal of controlled release : official journal of the Controlled Release Society 233, 101–113 (2016).

35. R. Guyon, S. Reinke, A. Truby, L. Sims, A. V. S. Hill, L. Bau, E. Stride, A. Milicic, Core-shell microcapsules compatible with routine injection enable prime/boost immunization against malaria with a single shot. Sci. Transl. Med. 17, eadw2256–eadw2256 (2025).

36. A. E. Witeof, N. M. Meinerz, K. D. Walker, H. H. Funke, R. L. Garcea, T. W. Randolph, A Single Dose, Thermostable, Trivalent Human Papillomavirus Vaccine Formulated Using Atomic Layer Deposition. J. Pharm. Sci. 112, 2223–2229 (2023).

37. A. E. Witeof, W. D. McClary, L. T. Rea, Q. Yang, M. M. Davis, H. H. Funke, C. E. Catalano, T. W. Randolph, Atomic-Layer Deposition Processes Applied to Phage λ and a Phage-like Particle Platform Yield Thermostable, Single-Shot Vaccines. J Pharm Sci 111, 1354–1362 (2022).

38. A. A. Cohen, J. R. Keeffe, A. V. Rorick, S. Rho, A.-C. P. Fils, L. Manasyan, H. Gao, P. N. P. Gnanapragasam, H. H. Funke, T. W. Randolph, R. L. Garcea, P. J. Bjorkman, Broad anti-sarbecovirus responses elicited by a single administration of mosaic-8 RBD-nanoparticle vaccine prepared using atomic layer deposition. iScience 28, 113649 (2025).

39. T. Randolph, H. Coleman, A. M. Rauch, H. Hubert, A. E. Witeof, K. D. Walker, H. H. Funke, R. L. Garcea, Superior immune responses from thermostable, single-administration rabies vaccines prepared using atomic layer deposition. J. Pharm. Sci. 114, 103936 (2025).

40. R. L. Garcea, N. M. Meinerz, M. Dong, H. Funke, S. Ghazvini, T. W. Randolph, Single-administration, thermostable human papillomavirus vaccines prepared with atomic layer deposition technology. Npj Vaccines 5, 45 (2020).

41. S. W. Brubaker, I. R. Walters, E. M. Hite, L. R. Antunez, E. L. Palm, H. H. Funke, B. L. Steadman, Demonstration of Tunable Control over a Delayed-Release Vaccine Using Atomic Layer Deposition. Vaccines 12, 761 (2024).

42. K. A. Strand, H. D’Angelo, A. M. Olson, I. R. Walters, E. M. Snyder, A. M. Ritter, E. Hite, Y. H. Wijesundara, A. B. Caplan, D. L. Ivanova, I. Allen, Y. Han, L. R. Antunez, A. K. Dey, B. L. Steadman, S. W. Brubaker, Atomic layering thermostable antigen and adjuvant (ALTA®) provides unique, controlled antigen release to improve immune response to vaccination. J. Control. Release 395, 114984 (2026).

43. D. L. Ivanova, M. S. Lewis, A. B. Caplan, I. R. Walters, E. M. Snyder, K. A. Strand, L. R. Antunez, S. Sengyee, S. B. Weiby, F. Urbano-Munoz, M. N. Burtnick, P. J. Brett, S. W. Brubaker, Subunit vaccination using Atomic Layering Thermostable Antigen and Adjuvant (ALTA®) platform elicits enhanced humoral and cellular immune responses. Front. Immunol. 17, 1787216 (2026).

44. D. M. King, X. Liang, A. W. Weimer, Functionalization of fine particles using atomic and molecular layer deposition. Powder Technol. 221, 13–25 (2012).

45. T. Atabey, R. W. Sanders, Y. Aldon, Germline-targeting Strategies to Induce bNAbs against HIV-1. Curr. HIV Res. 23 (2025), doi:10.2174/011570162x365229250527111140.

46. P. Broz, V. M. Dixit, Inflammasomes: mechanism of assembly, regulation and signalling. Nature Reviews Immunology 16, 407–420 (2016).

47. F. Lebre, C. H. Hearnden, E. C. Lavelle, Modulation of Immune Responses by Particulate Materials. Advanced Materials 28, 5525–5541 (2016).

48. Y. Li, Q. Jiang, Uncoupled pyroptosis and IL-1β secretion downstream of inflammasome signaling. Front. Immunol. 14, 1128358 (2023).

49. L. Hatscher, L. Amon, L. Heger, D. Dudziak, Inflammasomes in dendritic cells: Friend or foe? Immunol. Lett. 234, 16–32 (2021).

50. I. Zanoni, Y. Tan, M. D. Gioia, J. R. Springstead, J. C. Kagan, By Capturing Inflammatory Lipids Released from Dying Cells, the Receptor CD14 Induces Inflammasome-Dependent Phagocyte Hyperactivation. Immunity 47, 697–709.e3 (2017).

51. M. Idzko, H. Hammad, M. van Nimwegen, M. Kool, M. A. M. Willart, F. Muskens, H. C. Hoogsteden, W. Luttmann, D. Ferrari, F. D. Virgilio, J. C. Virchow, B. N. Lambrecht, Extracellular ATP triggers and maintains asthmatic airway inflammation by activating dendritic cells. Nat. Med. 13, 913–919 (2008).

52. F. Ghiringhelli, L. Apetoh, A. Tesniere, L. Aymeric, Y. Ma, C. Ortiz, K. Vermaelen, T. Panaretakis, G. Mignot, E. Ullrich, J.-L. Perfettini, F. Schlemmer, E. Tasdemir, M. Uhl, P. Génin, A. Civas, B. Ryffel, J. Kanellopoulos, J. Tschopp, F. André, R. Lidereau, N. M. McLaughlin, N. M. Haynes, M. J. Smyth, G. Kroemer, L. Zitvogel, Activation of the NLRP3 inflammasome in dendritic cells induces IL-1β–dependent adaptive immunity against tumors. Nat. Med. 15, 1170–1178 (2009).

53. M. Silva, Y. Kato, M. B. Melo, I. Phung, B. L. Freeman, Z. Li, K. Roh, J. W. V. Wijnbergen, H. Watkins, C. A. Enemuo, B. L. Hartwell, J. Y. H. Chang, S. Xiao, K. A. Rodrigues, K. M. Cirelli, N. Li, S. Haupt, A. Aung, B. Cossette, W. Abraham, S. Kataria, R. Bastidas, J. Bhiman, C. Linde, N. I. Bloom, B. Groschel, E. Georgeson, N. Phelps, A. Thomas, J. Bals, D. G. Carnathan, D. Lingwood, D. R. Burton, G. Alter, T. P. Padera, A. M. Belcher, W. R. Schief, G. Silvestri, R. M. Ruprecht, S. Crotty, D. J. Irvine, A particulate saponin/TLR agonist vaccine adjuvant alters lymph flow and modulates adaptive immunity. Sci. Immunol. 6, eabf1152 (2021).

54. P. Yousefpour, A. R. Ghosh, H. Chawla, R. Yeung, J. Gregory, K. Si, T. K. Remba, K. A. Rodrigues, M. B. Melo, J. Dye, J. M. Steichen, Y. Zhang, Y. Dong, M. Crispin, W. R. Schief, F. D. Batista, D. J. Irvine, Augmented germinal center and antibody responses to HIV Env trimers delivered as transmembrane immunogens by self-replicating RNA. Mol. Ther. 33, 4858–4873 (2025).

55. K. A. Rodrigues, C. A. Cottrell, J. M. Steichen, B. Groschel, W. Abraham, H. Suh, Y. Agarwal, K. Ni, J. Y. H. Chang, P. Yousefpour, M. B. Melo, W. R. Schief, D. J. Irvine, Optimization of an alum-anchored clinical HIV vaccine candidate. npj Vaccines 8, 117 (2023).

56. T. J. Moyer, Y. Kato, W. Abraham, J. Y. H. Chang, D. W. Kulp, N. Watson, H. L. Turner, S. Menis, R. K. Abbott, J. N. Bhiman, M. B. Melo, H. A. Simon, S. H.-D. la Mata, S. Liang, G. Seumois, Y. Agarwal, N. Li, D. R. Burton, A. B. Ward, W. R. Schief, S. Crotty, D. J. Irvine, Engineered immunogen binding to alum adjuvant enhances humoral immunity. Nat. Med. 26, 430–440 (2020).

57. T. Tokatlian, B. J. Read, C. A. Jones, D. W. Kulp, S. Menis, J. Y. H. Chang, J. M. Steichen, S. Kumari, J. D. Allen, E. L. Dane, A. Liguori, M. Sangesland, D. Lingwood, M. Crispin, W. R. Schief, D. J. Irvine, Innate immune recognition of glycans targets HIV nanoparticle immunogens to germinal centers. Science 363, 649–654 (2019).

58. S. H. Bhagchandani, L. Yang, J. H. Lam, L. Maiorino, E. Ben-Akiva, K. A. Rodrigues, A. Romanov, H. Suh, A. Aung, S. Wu, A. Wadhera, A. K. Chakraborty, D. J. Irvine, Two-dose priming immunization amplifies humoral immunity by synchronizing vaccine delivery with the germinal center response. Sci. Immunol. 9, eadl3755–eadl3755 (2024).

59. H. H. Tam, M. B. Melo, M. Kang, J. M. Pelet, V. M. Ruda, M. H. Foley, J. K. Hu, S. Kumari, J. Crampton, A. D. Baldeon, R. W. Sanders, J. P. Moore, S. Crotty, R. Langer, D. G. Anderson, A. K. Chakraborty, D. J. Irvine, Sustained antigen availability during germinal center initiation enhances antibody responses to vaccination. Proc National Acad Sci 113, E6639–E6648 (2016).

60. K. A. Rodrigues, Y. J. Zhang, J. Lam, A. Aung, D. M. Morgan, A. Romanov, L. Maiorino, P. Yousefpour, G. Gibson, G. Ozorowski, J. R. Gregory, P. Amlashi, R. Van, M. Buckley, A. B. Ward, W. R. Schief, J. C. Love, D. J. Irvine, Vaccines combining slow release and follicle targeting of antigens increase germinal center B cell diversity and clonal expansion. Sci. Transl. Med. 17, eadw7499 (2025).

61. C. Havenar-Daughton, J. H. Lee, S. Crotty, Tfh cells and HIV bnAbs, an immunodominance model of the HIV neutralizing antibody generation problem. Immunol Rev 275, 49–61 (2017).

62. M. Pauthner, C. Havenar-Daughton, D. Sok, J. P. Nkolola, R. Bastidas, A. V. Boopathy, D. G. Carnathan, A. Chandrashekar, K. M. Cirelli, C. A. Cottrell, A. M. Eroshkin, J. Guenaga, K. Kaushik, D. W. Kulp, J. Liu, L. E. McCoy, A. L. Oom, G. Ozorowski, K. W. Post, S. K. Sharma, J. M. Steichen, S. W. de Taeye, T. Tokatlian, A. T. de la Peña, S. T. Butera, C. C. LaBranche, D. C. Montefiori, G. Silvestri, I. A. Wilson, D. J. Irvine, R. W. Sanders, W. R. Schief, A. B. Ward, R. T. Wyatt, D. H. Barouch, S. Crotty, D. R. Burton, Elicitation of Robust Tier 2 Neutralizing Antibody Responses in Nonhuman Primates by HIV Envelope Trimer Immunization Using Optimized Approaches. Immunity 46, 1073–1088.e6 (2017).

63. K. A. Rodrigues, S. A. Rodriguez-Aponte, N. C. Dalvie, J. H. Lee, W. Abraham, D. G. Carnathan, L. E. Jimenez, J. T. Ngo, J. Y. H. Chang, Z. Zhang, J. Yu, A. Chang, C. Nakao, B. Goodwin, C. A. Naranjo, L. Zhang, M. Silva, D. H. Barouch, G. Silvestri, S. Crotty, J. C. Love, D. J. Irvine, Phosphate-mediated coanchoring of RBD immunogens and molecular adjuvants to alum potentiates humoral immunity against SARS-CoV-2. Sci. Adv. 7, eabj6538 (2021).

64. H. J. Coleman, A. Rauch, E. Langsfeld, K. Anderson, U. Parlikar, H. H. Funke, R. L. Garcea, T. W. Randolph, Lipid-free, thermostable mRNA vaccines prepared using atomic layer deposition. J. Pharm. Sci. 115, 104066 (2026).

65. C. A. Schneider, W. S. Rasband, K. W. Eliceiri, NIH Image to ImageJ: 25 years of image analysis. Nat. Methods 9, 671–675 (2012).

66. J. Schindelin, I. Arganda-Carreras, E. Frise, V. Kaynig, M. Longair, T. Pietzsch, S. Preibisch, C. Rueden, S. Saalfeld, B. Schmid, J.-Y. Tinevez, D. J. White, V. Hartenstein, K. Eliceiri, P. Tomancak, A. Cardona, Fiji: an open-source platform for biological-image analysis. Nat. Methods 9, 676–682 (2012).

