## Supplementary Information for "Atomic layer deposition for core-shell microparticle vaccines enabling programmable antigen delivery and enhanced humoral immune responses"

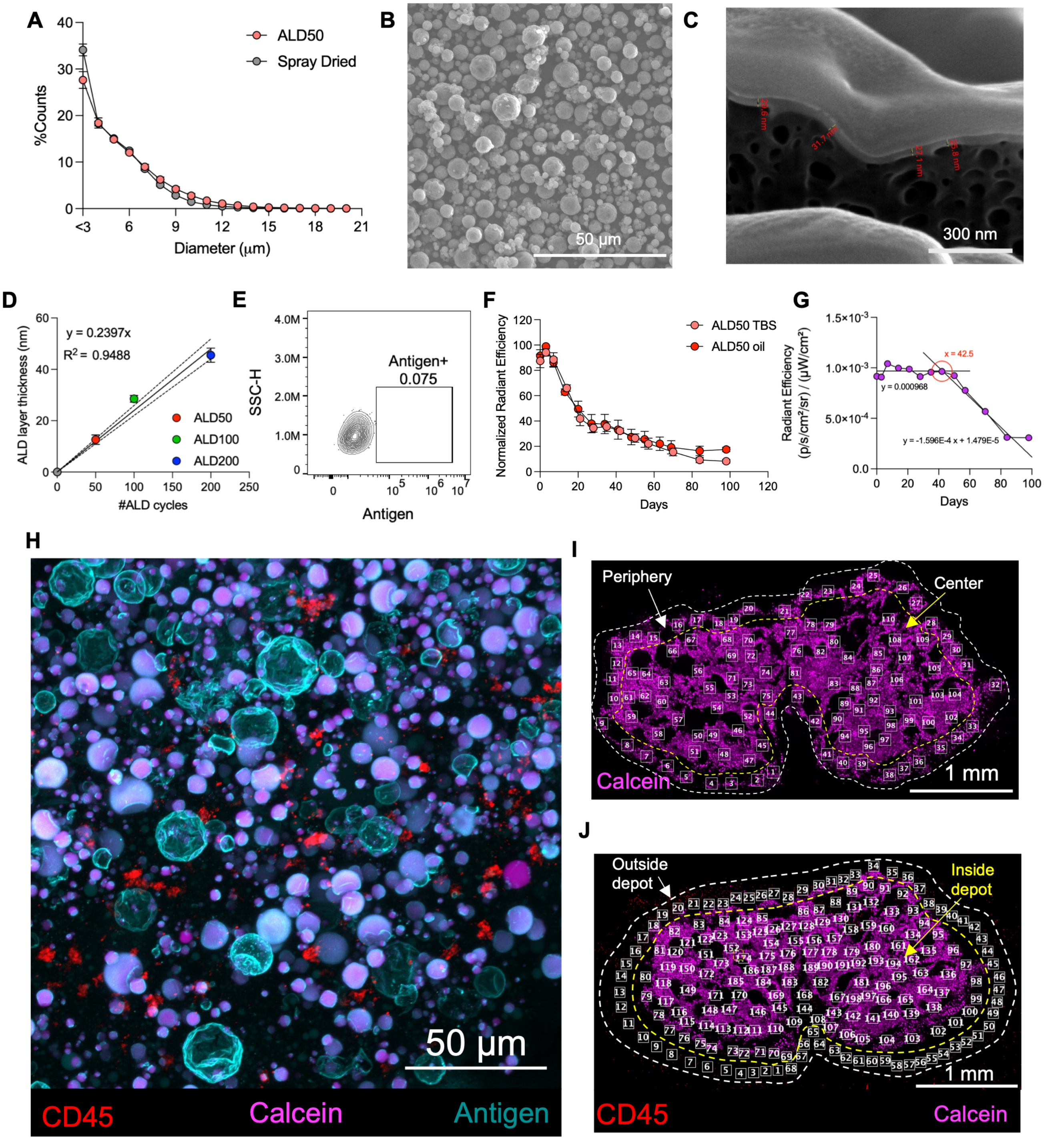


**Fig. S1. Physical characterization of ALD microparticles and functional behavior at injection sites.** **A)** Antigen was spray-dried using formulation 1 and coated with 50 layers of alumina. Particle size distributions were determined for spray-dried and ALD microparticles by flow imaging microscopy. **B)** Morphology of spray-dried particles was visualized via scanning electron microscopy. **C-D)** Thickness of alumina layers deposited by ALD on microparticles was determined by FIB-SEM. Shown are a representative FIB-SEM image of the alumina shell in ALD100 particles **(C)** and measured alumina shell thickness as a function ALD cycle number **(D)**. Each ALD cycle deposited 0.24±0.02 nm alumina, as determined from a linear plot ($r^2$=0.95) of alumina layer thickness vs. number of ALD cycles. **E)** BMDCs (*n* = 4 samples/group) were incubated with 5 µg/mL free Alexa Fluor 647-labeled antigen for 24 hr, followed by flow cytometry analysis. Shown is representative flow cytometry plot of antigen uptake. **F)** Albino C57BL/6 mice (*n* = 5 animals/group) were injected s.c. with ALD50 particles encapsulating 10 µg Alexa Fluor 647–labeled antigen suspended in either triglyceride oil or Tris-buffered saline. Antigen persistence at the injection site was monitored longitudinally by whole-animal fluorescence imaging and quantified as normalized radiant efficiency over time. **G)** Animals were immunized with ALD particles encapsulating fluorescent antigen as in **Fig. 3B** and antigen fluorescence at the injection site was tracked over time. To estimate the time point when the bulk of antigen clearance begins, the initial plateau and subsequent decay regions of the fluorescence data were fit by linear regression, and the time point for intersection of the two best fit lines was taken as the time of release initiation. **H-J)** C57BL/6 mice (*n* = 4 animals/group) were immunized with 5 µg Alexa Fluor 647-labeled antigen and 7.5 µg calcein administered as ALD particles suspended in oil. Injection sites were excised and imaged by fluorescence microscopy to assess spatial distribution of particle integrity and immune cell infiltration within the depot. **H)** Shown are representative high magnification images of the periphery at day 14. **I)** Representative low-magnification fluorescence image of an ALD depot (calcein, magenta). ROIs were manually defined across two spatial zones, periphery (outer rim, white dashed boundary) and center (yellow dashed boundary), and the fraction of intact particles in each zone was quantified using Mander's coefficient. **J)** A similar approach was applied to quantify immune cell (CD45, red) distribution, with ROIs defined outside (between white and yellow dashed lines) and inside (within yellow dashed line) the depot. Immune cell intensity inside the depot was normalized to the mean outside intensity to yield a relative infiltration measure.

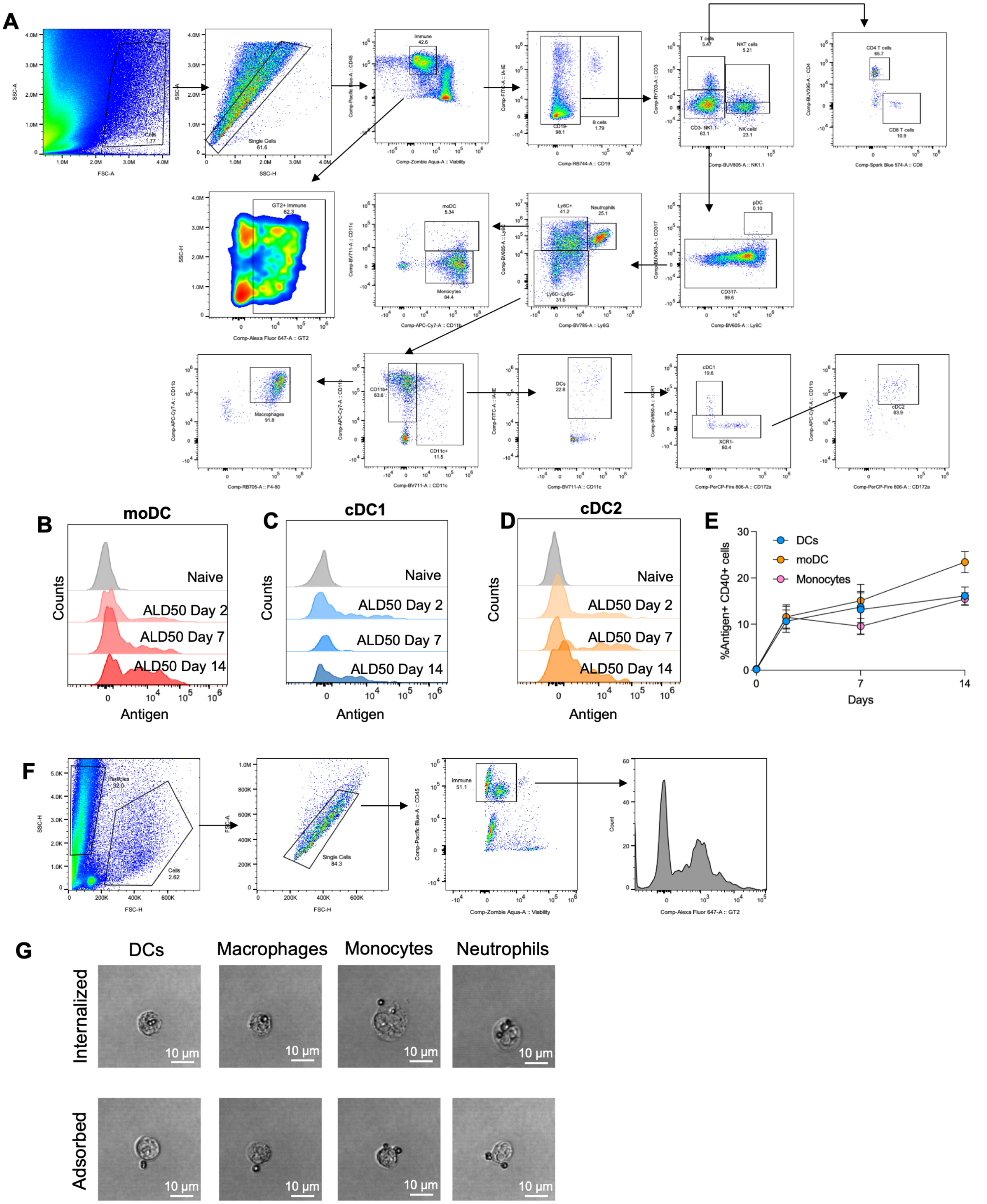


**Fig. S2. Flow-cytometric identification of immune cell populations and antigen uptake at ALD injection site. A)** Representative flow-cytometry gating strategy used to identify major immune cell populations from skin tissue following subcutaneous immunization. After exclusion of debris, doublets, and dead cells, $CD45^+$ leukocytes were subdivided as follows: B cells ($CD19^+$); T cells ($CD19^-CD3^+$), with $CD4^+$ and $CD8^+$ T cells identified within the T-cell gate; NK cells ($CD19^-CD3^-NK1.1^+$); NKT cells ($CD19^-CD3^+NK1.1^+$); plasmacytoid dendritic cells (pDCs; $CD19^-CD3^-CD317^+$); neutrophils ($CD19^-CD3^-CD317^-Ly6C^+Ly6G^+$); monocytes ($CD19^-CD3^-CD317^-Ly6C^+Ly6G^-CD11b^+CD11c^-$); monocyte-derived dendritic cells (moDCs; $CD19^-CD3^-CD317^-Ly6C^+Ly6G^-CD11b^+CD11c^+$); macrophages ($CD19^-CD3^-CD317^-Ly6C^-Ly6G^-CD11b^+CD11c^-F4/80^+$); and dendritic cells (DCs; $CD19^-CD3^-CD317^-Ly6C^-Ly6G^-CD11c^+I\text{-}A/I\text{-}E^+$). Conventional DC subsets were further defined as cDC1 ($XCR1^+$) and cDC2 ($XCR1^-CD172a^+CD11b^+$). **B-D)** Representative histograms showing uptake of Alexa Fluor 647–labeled antigen by moDCs **(B)**, cDC1 **(C)**, and cDC2 **(D)** at the injection site at baseline (naïve) and at days 2, 7, and 14 following immunization with ALD particles. **(E)** Frequency of $antigen^+$ activated DCs over time. **F)** Imaging cytometry gating for immune cells at the injection site, gating on FSC/SSC, followed by gating for single cells, live/CD45+ cells, and antigen fluorescence signal. **G)** Representative cytometer images of myeloid cell types showing internalized or adsorbed ALD particles from the injection site.

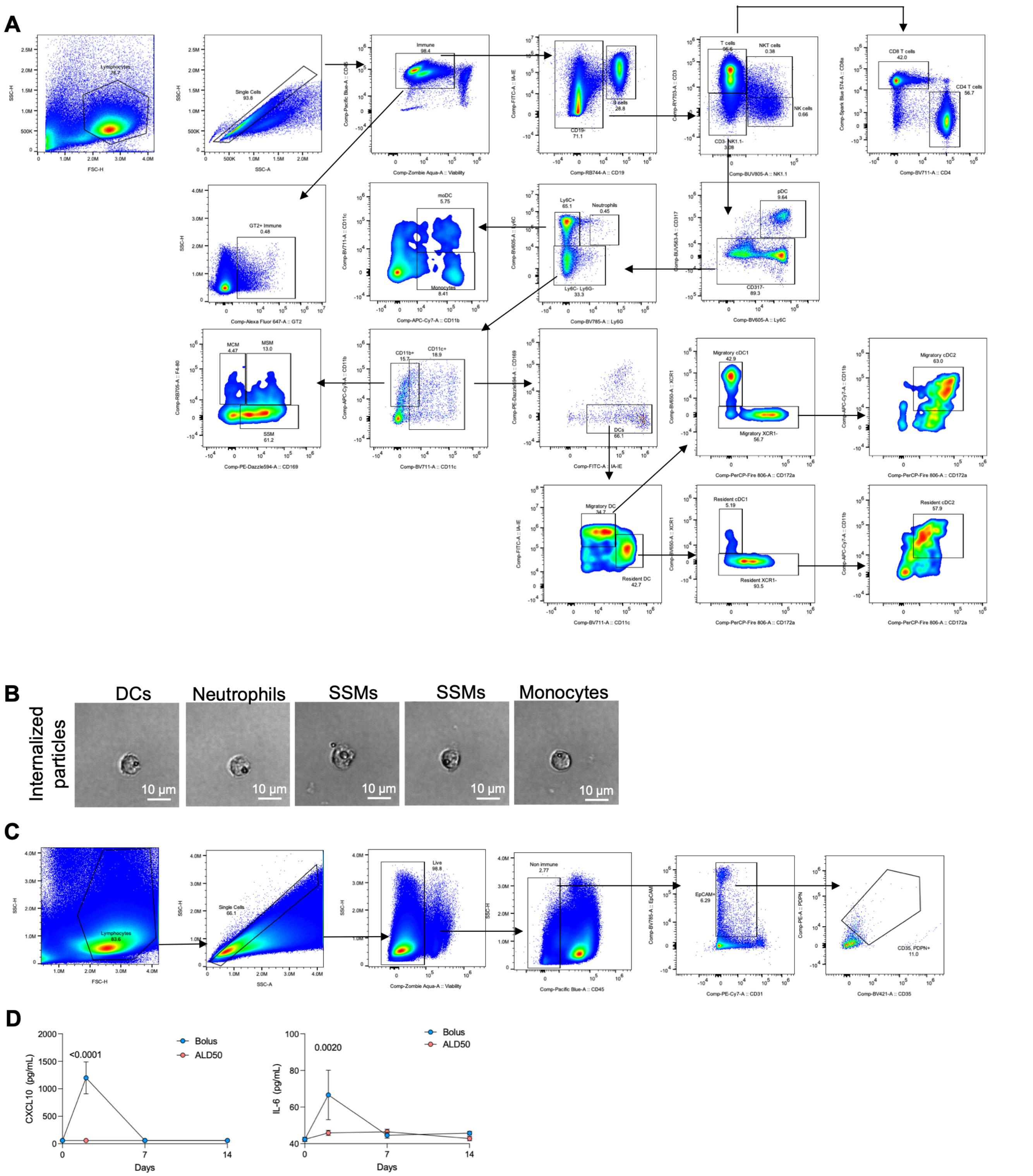


**Fig. S3. Flow-cytometric identification of lymph node immune cells. A)** Flow cytometry gating strategy used to identify immune cell populations in draining lymph nodes following subcutaneous immunization. After exclusion of debris, doublets, and dead cells, $CD45^{+}$ leukocytes were subdivided as follows: B cells ($CD19^{+}$); T cells ($CD19^{-}CD3^{+}$), with $CD4^{+}$ and $CD8^{+}$ T cells identified within the T-cell gate; NK cells (CD19$^-$CD3$^-$NK1.1$^+$); NKT cells (CD19$^-$CD3$^+$NK1.1$^+$); plasmacytoid dendritic cells (pDCs; CD19$^-$CD3$^-$CD317$^+$); neutrophils (CD19$^-$CD3$^-$CD317$^-$Ly6C$^+$Ly6G$^+$); monocytes (CD19$^-$CD3$^-$CD317$^-$Ly6C$^+$Ly6G$^-$CD11b$^+$CD11c$^-$); and monocyte-derived dendritic cells (moDCs; CD19$^-$CD3$^-$CD317$^-$Ly6C$^+$Ly6G$^-$CD11b$^+$CD11c$^+$). Macrophages were defined within the CD19$^-$CD3$^-$CD317$^-$Ly6C$^-$Ly6G$^-$CD11b$^+$CD11c$^{lo}$ gate and subdivided into subcapsular sinus macrophages (SSMs; CD169$^+$F4/80$^-$), medullary sinus macrophages (MSMs; CD169$^+$F4/80$^+$), and medullary cord macrophages (MCMs; CD169$^-$F4/80$^+$). Dendritic cells (DCs) were defined within the CD19$^-$CD3$^-$CD317$^-$Ly6C$^-$Ly6G$^-$CD11c$^+$I-A/I-E$^+$ gate and further restricted to CD169$^-$ cells (I-A/I-E$^+$CD169$^-$) to exclude CD169$^+$ macrophages. DCs were separated into migratory DCs (CD11c$^{lo}$ I-A/I-E$^{hi}$) and lymph node–resident DCs (CD11c$^{hi}$ I-A/I-E$^{lo}$). Migratory and resident DC subsets were each further classified into cDC1 (XCR1$^+$) and cDC2 (XCR1$^-$CD172a$^+$CD11b$^+$) populations. **B)** Representative cytometer images of myeloid cell populations showing internalized ALD particles in the lymph nodes. **C)** Flow cytometry gating strategy used to identify follicular dendritic cells (FDCs) in draining lymph nodes and to quantify antigen retention. After exclusion of debris, doublets, and dead cells, FDCs were defined as CD45$^-$EpCAM$^+$CD31$^-$CD35$^+$PDPN$^+$ cells. **D)** C57BL/6 mice (*n*=5 animals/group) were immunized with 10 µg antigen delivered either as a liquid bolus dose with 5 µg SMNP adjuvant or encapsulated within ALD50 particles suspended in triglyceride oil. Serum cytokines and chemokines were measured by Legendplex. Error bars represent s.e.m. Statistical significance was determined by two-way ANOVA with Šidák's multiple-comparison test **(D).**

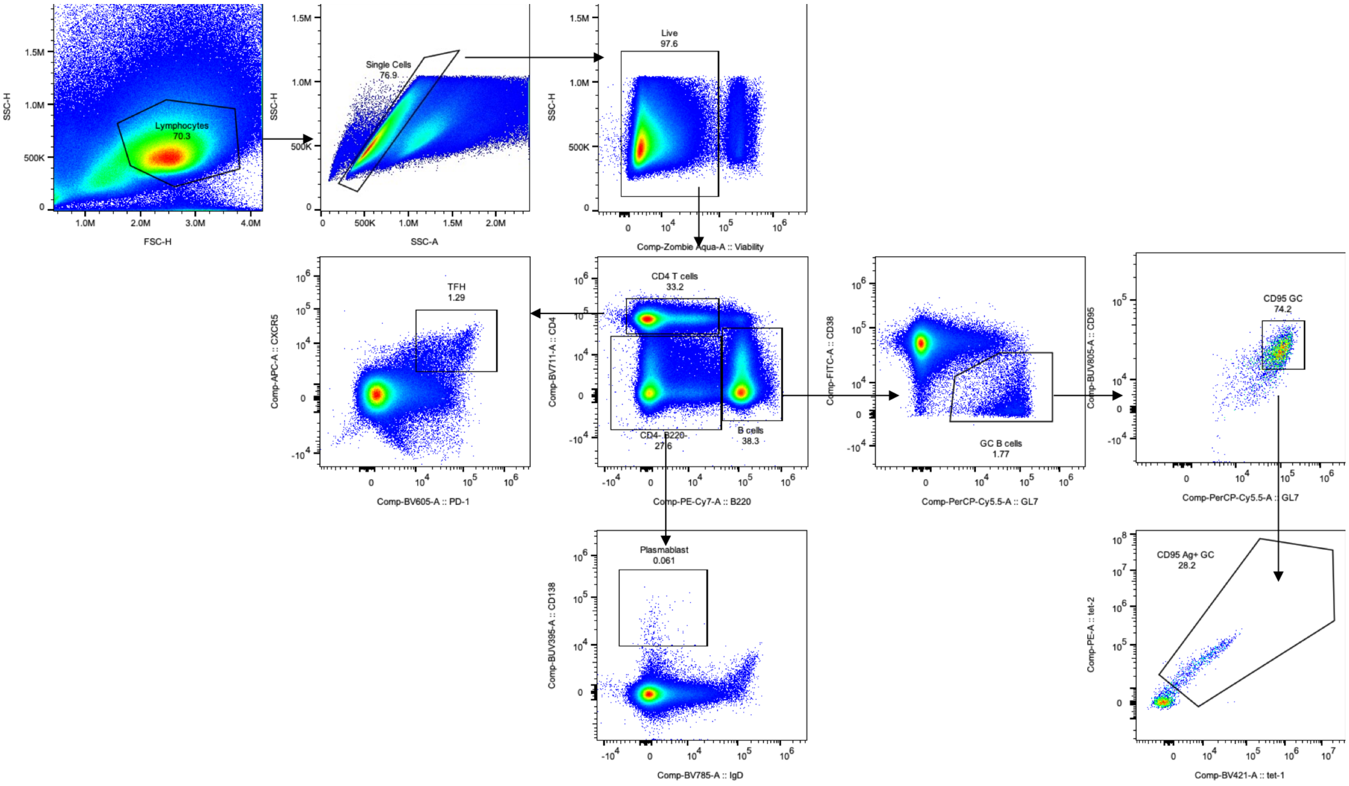


**Fig. S4. Flow -cytometric identification of follicular dendritic cell antigen retention and germinal center responses in draining lymph nodes.** Flow cytometry gating strategy used to quantify germinal center responses in lymph nodes. Germinal center (GC) B cells were defined as CD4$^-$B220$^+$GL7$^+$CD38$^{lo}$CD95$^+$ cells. Antigen-specific GC B cells were identified within the GC B-cell gate by dual binding to antigen tetramers labeled with BV421 and PE. T follicular helper ($T_{FH}$) cells were defined as B220$^-$CD4$^+$PD-1$^+$CXCR5$^+$ cells. Plasmablasts were defined as B220$^-$CD4$^-$IgD$^-$CD138$^+$ cells.

**Supplementary Table 1. Spray drying formulations evaluated for HIV Env trimer encapsulation.**

| Formulation | Histidine (mM) | Tween 20 (wt%) | Hydroxyethylstarch (wt%) | pH |
|---|---|---|---|---|
| F1 | 50 | 0.2 | 2.5 | 6.5 |
| F2 | 50 | 0.2 | 0 | 6.5 |
| F3 | 50 | 0 | 2.5 | 6.5 |
| F4 | 50 | 0 | 0 | 6.5 |
| F5 | 50 | 0.2 | 2.5 | 5.5 |
| F6 | 50 | 0.2 | 0 | 5.5 |
| F7 | 50 | 0 | 2.5 | 5.5 |
| F8 | 50 | 0 | 0 | 5.5 |
| F9 | 10 | 0.2 | 2.5 | 6.5 |
| F10 | 10 | 0.2 | 0 | 6.5 |
| F11 | 10 | 0 | 2.5 | 6.5 |
| F12 | 10 | 0 | 0 | 6.5 |
| F13 | 10 | 0.2 | 2.5 | 5.5 |
| F14 | 10 | 0.2 | 0 | 5.5 |
| F15 | 10 | 0 | 2.5 | 5.5 |
| F16 | 10 | 0 | 0 | 5.5 |

All samples contained 50 µg/mL N332-GT2 trimer, 40 mM ammonium acetate, and 9.5 wt% trehalose.
